# Protonation/deprotonation-driven switch for the redox stability of low-potential [4Fe-4S] ferredoxin

**DOI:** 10.1101/2024.09.12.612615

**Authors:** Kei Wada, Kenji Kobayashi, Iori Era, Yusuke Isobe, Taigo Kamimura, Masaki Marukawa, Takayuki Nagae, Kazuki Honjo, Noriko Kaseda, Yumiko Motoyama, Kengo Inoue, Masakazu Sugishima, Katsuhiro Kusaka, Naomine Yano, Keiichi Fukuyama, Masaki Mishima, Yasutaka Kitagawa, Masaki Unno

## Abstract

Ferredoxin is a small iron-sulfur protein and acts as an electron carrier. Low-potential ferredoxins harbor [4Fe-4S] cluster(s), which play(s) a crucial role as the redox center. Low-potential ferredoxins are able to cover a wide range of redox potentials (−700 to −200 mV); however, the mechanisms underlying the factors which control the redox potential are still enigmatic. Here, we determined the neutron structure of ferredoxin from *Bacillus thermoproteolyticus*, and experimentally revealed the exact hydrogen-bonding network involving the [4Fe-4S] cluster. The density functional theory calculations based on the hydrogen-bonding network revealed that protonation states of the sidechain of Asp64 close to the [4Fe-4S] cluster critically affected the stability of the reduced state in the cluster. These findings provide the first identification of the intrinsic control factor of redox potential for the [4Fe-4S] cluster in low-potential ferredoxins.

## Introduction

Ferredoxin is known as a simple iron-sulfur (Fe-S) protein, and is ubiquitous in plants, animals, and prokaryotes(1, 2). Ferredoxins function as electron carriers for fundamental metabolic processes such as gluconeogenesis/glycolysis, photosynthesis, nitrogen fixation, and assimilation of hydrogens and sulfur species. Therefore, ferredoxins have a variety of redox partners. In photosynthetic organisms, ferredoxin receives an electron from photosystem I complex and serves as the electron donor for the reduction of NADP^+^ (ferredoxin:NADP^+^ oxidoreductase), pyruvate (pyruvate:ferredoxin oxidoreductase), nitrate (nitrate reductase), and sulfite (sulfite reductase) to generate sources of energy(3). In bacteria, oxidized ferredoxins are reduced *via* a reverse reaction such as by bacterial type ferredoxin:NADP^+^ oxidoreductase and pyruvate:ferredoxin oxidoreductase, and electrons in the reduced ferredoxin are utilized by ferredoxin-dependent enzymes in various metabolism cascades(4).

Typical bacterial ferredoxin is a small protein consisting of approximately one hundred amino acid residues, primarily (approximately 20%) negatively charged residues, with an isoelectric point (pI) that is usually below pH 4.0. The ferredoxin also has four conserved cysteine residues to which the Fe-S cluster is ligated in most cases. The types of the cluster are known as the [4Fe-4S] types(5) (Fig.1). [4Fe-4S] ferredoxins may be further classified into low-potential and high-potential ferredoxins. Intriguingly, [4Fe-4S] ferredoxins exhibit a wide range of redox potentials while having the same form of the cluster in the active site(6). For instance, low-potential ferredoxins harboring [4Fe-4S] exhibit redox potentials typically ranging from −700 mV to −200 mV; in contrast, high-potential ferredoxins have redox potentials of +100 mV to +360 mV(7). Low-potential [4Fe-4S] ferredoxins, which are typical bacterial-type ferredoxins, reduce most of the redox partners at physiological potentials, whereas high-potential ferredoxins called HiPIPs usually accept electrons from most redox partners. Notably, electron transfer events by low-potential [4Fe-4S] ferredoxins are remarkable reactions; the low-potential ferredoxin receives an electron and must maintain the reduced [4Fe-4S] cluster until it transfers the electron to the redox partner proteins. How the low-potential [4Fe-4S] cluster maintains stability in the reduced form is unknown. The surrounding environment of the [4Fe-4S] cluster must be conferred and controls electron competency/stability; however, the mechanism underlying the sophisticated regulation of the redox potential and electron flow are still elusive.

**Fig. 1.**
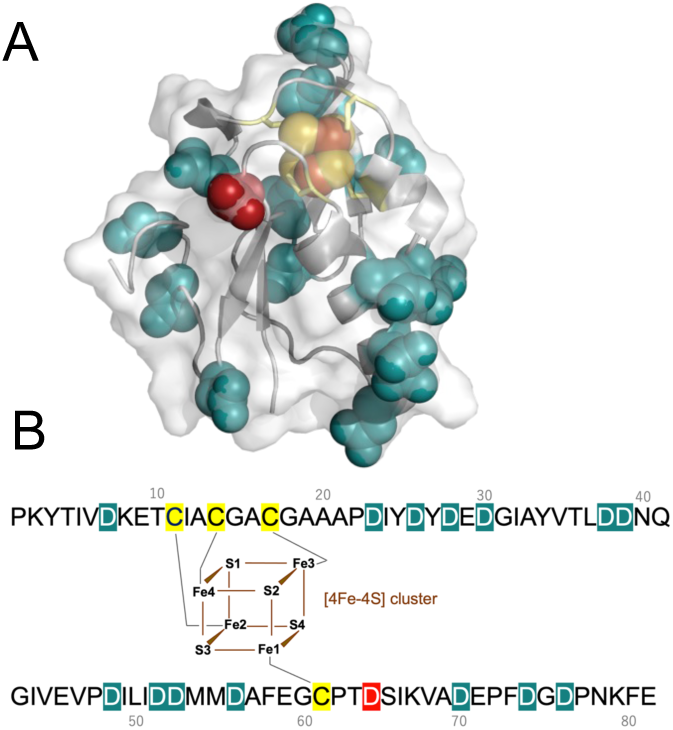
Ternary and primary structures of low-potential [4Fe-4S] *Bt*Fd. (*A*) The 3D structure of BtFd (PDB id: 1IQZ). The [4Fe-4S] cluster and aspartate residues are shown as a space-filling model, and the ligand cysteines for the Fe-S cluster are shown as yellow sticks. The surface drawing is superposed on the cartoon model. (*B*) Whole amino acid sequence of BtFd. The ligating manner of the [4Fe-4S] cluster is illustrated, and the ligand cysteine and aspartate residues are highlighted.

Computer simulations using model complexes of the Fe-S cluster showed that the covalent nature of the Fe-S cluster decreases due to hydrogen bonds with the surrounding amino acid residues, and the redox potential correlates well with the covalent nature(8). This indicates that the redox potential may be controlled by the hydrogen bonds between the main chain amide nitrogen and the sulfur in iron-sulfur clusters (NH…S). In contrast, Solomon and co-workers demonstrated the substantial influence of hydration, supplied by water molecules, on the variation in reactivity by X-ray absorption spectroscopy(9). The Fe-S covalency is much lower in natively hydrated [4Fe-4S] ferredoxin active sites but increases upon water removal. The density functional theory (DFT) calculations have supported a correlation between Fe-S covalency and ease of oxidation and therefore suggest that hydration accounts for most of the differences between ferredoxin redox potentials. However, the most critical problem in the computational calculations for the protein is that artificial coordinates for hydrogen atoms, which are predicted and generated computationally, have been used thus far for the calculations.

The precise locations of the hydrogen atoms in proteins are difficult to determine experimentally because this requires high-resolution structural information. So far, the exceptional ultra-high-resolution structure at 0.48 Å for HiPIP from *Thermochromatium tepidum* has been reported by X-ray crystallography(10), and the details of the hydrogen bonding networks surrounding the [4Fe-4S] cluster were determined. Using the charge-density analysis focused the valence electron, and the iron 3*d* and sulfur 3*p* electrons of the high-potential (positivity potential) [4Fe-4S] cluster were visualized for the first time. This is currently the most accurate analysis of the protein structure, but few indications regarding the stability of the redox state of the [4Fe-4S] cluster were obtained. More recently, the neutron structure of the HiPIP has been reported, indicating that the orientation of the amide proton of Cys75, one of the ligand residues for the Fe-S cluster, was probably altered in the reduced and oxidized states, possibly because of the electron storage capacity of the iron-sulfur cluster(11).

The high-resolution structure of the low-potential [4Fe-4S] ferredoxin was reported using a gram-positive bacterium *Bacillus thermoproteolyticus* (*Bt*Fd). *Bt*Fd is a typical bacterial ferredoxin and is a small protein, a protein with pI in the acidic region, that consists of 81 residues and one [4Fe-4S] cluster (Fig. 1). The X-ray structure of *Bt*Fd has been reported at a 0.92 Å resolution(12), but the hydrogen atoms, particularly around the Fe-S clusters, could not be assigned on the electron density map. The DFT calculations, based on the *Bt*Fd structure, reported that the Fe-S covalencies of [4Fe-4S] clusters differ considerably from HiPIP because of hydrogen bonds with water molecules(9). In this case, the hydrogen-bonding networks surrounding the Fe-S clusters also used the putative positions based on the calculations.

Neutron crystallography is a powerful analytic method because the hydrogen atoms can be visualized and the radiation damage is negligible. In this study, we proceeded to experimentally visualize the hydrogen bonds around the oxidized [4Fe-4S] cluster in *Bt*Fd by neutron crystal structure analysis under ambient air conditions and using the DFT method based on the experimentally determined hydrogen bonding network. The DFT calculations revealed that the protonation states of Asp64 located close to the reduced/oxidized [4Fe-4S] cluster significantly affected the redox stability of the cluster. Furthermore, the autooxidation kinetics of the reduced [4Fe-4S] cluster of *Bt*Fd prepared under anaerobic conditions supported the notion that Asp64 is critically involved in the stability of the reduced [4Fe-4S] cluster. These findings provide the first identification of intrinsic factors controlling the redox stability of the [4Fe-4S] cluster in low-potential ferredoxins.

## Results

### Overall neutron structure of the low-potential [4Fe-4S] ferredoxin

We have successfully determined the crystallization conditions of *Bt*Fd in which we could obtain large crystals (>2 mm^3^) using the dialysis method. The neutron structure of *Bt*Fd at room temperature was refined to a 1.60 Å resolution. The *R*-factor and free *R*-factor were 16.7% and 18.4%, respectively (Table 1). A total of 819 H/D atoms and 60 hydrating water molecules were identified (Fig. 2A, Table 1, and Fig. S1). The overall structure was nearly identical to the previously reported cryogenic X-ray structure (PDB ID: 1IQG) at 0.92 Å resolution, with a rms deviation of 0.16 Å for Cα atoms when all residues (81 residues) were superimposed with least-squares fitting. Although the neutron-scattering length density map clearly shows the [4Fe-4S] cluster, the density for sulfur atoms was very low compared with most of the other atoms including the iron atoms (Fig. 2B). This is a reasonable result because the sulfur atom has a smaller neutron-scattering length *b*_c_ (10^-12^ cm) (^32^S=0.2804) than other atoms (^56^Fe=0.994, ^12^C=0.6651, ^14^N=0.937, ^16^O=0.5803). The ability to clearly distinguish between iron atoms and sulfur atoms is one of the characteristics of neutron structure analysis. We also collected the X-ray diffraction data from the same crystal used for neutron diffraction intensity measurements, and the 1.45 Å resolution X-ray structure, obtained at room temperature, was also refined (Table 1). The electron density map of the [4Fe-4S] cluster showed the respective locations of iron and sulfur atoms with strong electron densities (Fig. 2C). Consequently, the joint refinement of the neutron structure with the X-ray structure gave the precise coordinates of the [4Fe-4S] cluster.

**Fig. 2.**
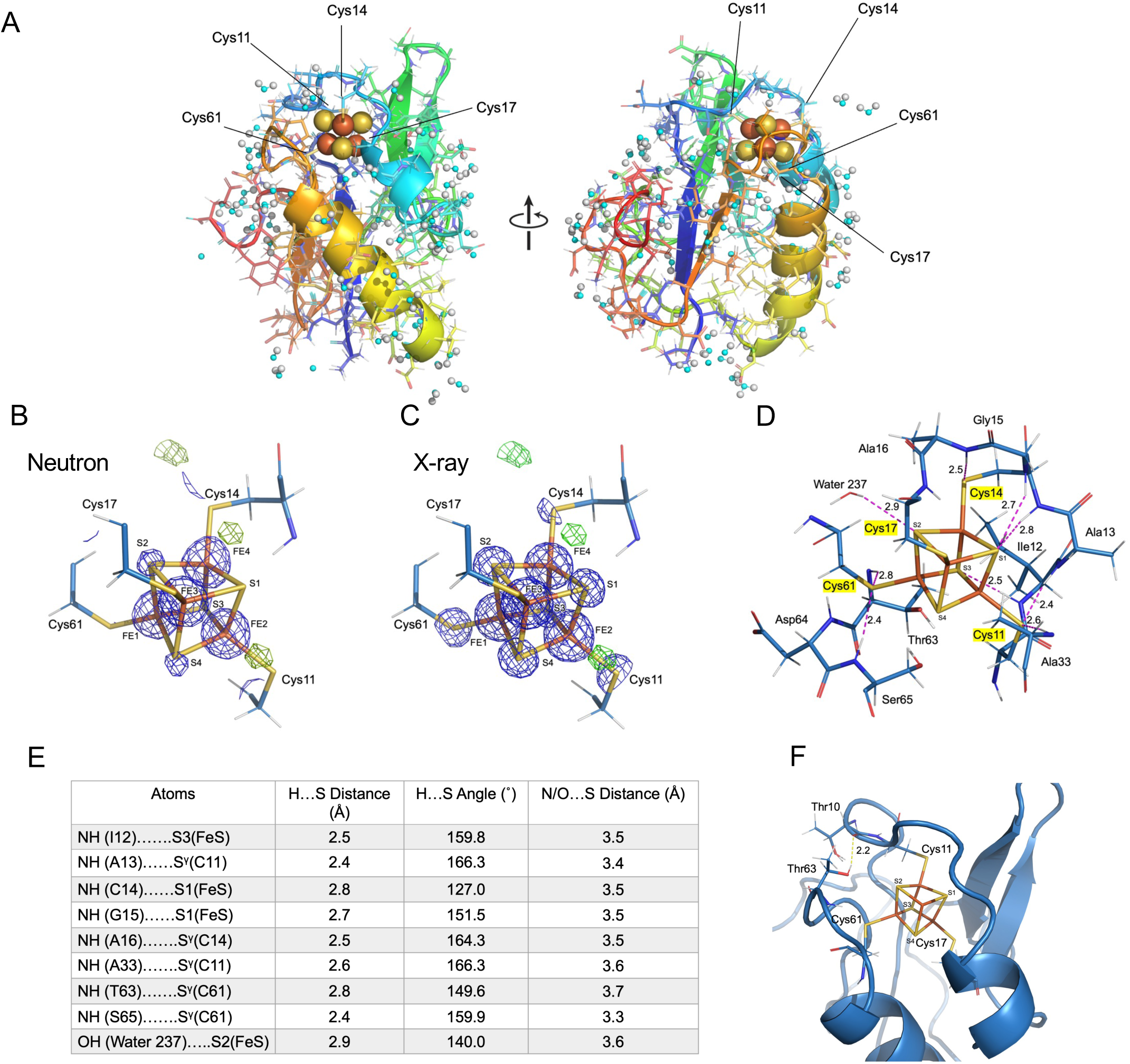
Neutron structure of low-potential [4Fe-4S] *Bt*Fd. (*A*) Overall neutron structure of *Bt*Fd. *Bt*Fd polypeptides are colored in rainbow colors from blue (N-terminus) to red (C-terminus). The [4Fe-4S] cluster and water molecules are shown as a space-filling model. White and cyan balls indicate hydrogen (deuterium) and oxygen atoms, respectively. The figure on the right is the left one rotated approximately 90°. (*B*) The 2*F*_o_–*F*_c_ (1.0 sigma: blue cage) and *F*_o_–*F*_c_ (3.0 sigma: green cage) neutron-scattering length density map around the [4Fe-4S] cluster and the structural model. The 2*F*_o_–*F*_c_ map is shown within 1 Å around the [4Fe-4S] cluster for clarity. (*C*) X-ray electron density map of the corresponding region of (*B)*. 2*F*_o_–*F*_c_ (5.0 sigma) and *F*_o_–*F*_c_ (3.0 sigma) maps are shown in blue and green, respectively. (*D*) Hydrogen bond with the [4Fe-4S] cluster and its ligand Sγ atoms of cysteines. The hydrogen (deuterium), oxygen, carbon, and nitrogen atoms are shown in white, red, cyan, and blue, respectively. The hydrogen bonds indicate pink broken lines and the distance between the amide hydrogen and sulfur atoms is shown in angstroms. (*E*) The distances and angles of the hydrogen bonds. The distance/angle between the hydrogen and the sulfur are shown on the left and the center columns, and the distance between nitrogen/oxygen and sulfur are shown on the right column. (*F*) Hydrogen bond between Thr63-OH and the main chain –CO of Thr10.

**Table 1.** Statistics for Neutron and X-ray Diffraction Data.

| Data Collection |  |  |
| --- | --- | --- |
|  | Neutron | X-ray |
| Beamline | iBIX at MLF in J-PARC | NW12A at PF-AR |
| Wavelength (Å) | 1.5 – 4.5 | 1.0000 |
| Exposure time | 6 hours/1 set | 0.5 second/0.5° |
| Number of datasets (neutron) or images (X-ray) | 58 (15 days) | 720 |
| Transmittance (%) | 100 | 5 |
| Resolution (Å) | 15.12 – 1.60<br>(1.69 – 1.60) | 50.00 – 1.45<br>(1.48 – 1.45) |
| Space group | C2 |  |
| Cell parameters (Å, °) | $a = 70.5, b = 37.9, c = 33.0$<br>$\beta = 106.0$ | |
| Multiplicity | 7.5 (6.2) | 6.7 (6.1) |
| Completeness (%) | 94.2 (76.8) | 100 (100) |
| $R_{\text{merge}}$ | 0.137 (0.369) | 0.080 (0.553) |
| $I/\sigma(I)$ | 12.9 (4.9) | 25.2 (2.5) |
| Joint refinement |  |  |
| $R_{\text{work}}/R_{\text{free}}$ | 0.167/0.184 | 0.147/0.159 |
| *r.m.s Bond lengths (Å)* | 0.010 | – |
| *r.m.s Bond angles (°)* | 1.067 | – |
| *Total No. of hydrogen atom* | 598 | – |
| *Total No. of deuterium atom* | 221 | – |
| *No. of H atoms in protein* | 598 | – |
| *No. of D atoms in protein* | 110 | – |
| *No. of H atoms in water* | 0 | – |
| *No. of D atoms in water* | 111 | – |

### Hydrogen atoms around the low-potential [4Fe-4S] cluster

In the neutron structure, the hydrogen atoms around the [4Fe-4S] cluster were visualized, and the exact direction of hydrogen bonds could be determined (Fig. 2D and E). The Sγ of cysteine residues, as ligands of the [4Fe-4S] cluster, formed five hydrogen bonds with the main chain amino groups; Cys11-Sγ formed two hydrogen bonds with Ala13-NH and Ala33-NH, Cys14-Sγ bonded to Ala16-NH, and Cys61-Sγ bonded to Ile12-NH and Ser65-NH. In contrast, the Sγ of Cys17 formed no hydrogen bond. The sulfur in the [4Fe-4S] cluster also formed hydrogen bonds; [4Fe-4S]-S1 with Gly15-NH and [4Fe-4S]-S3 with Ile12-NH. In addition, we observed the hydrogen atom of the water molecule (Water 237) close to the [4Fe-4S] cluster, which was bonded to [4Fe-4S]-S1 atom. In total, 9 hydrogen bonds were clearly identified around the [4Fe-4S] cluster (Fig. 2E). Nevertheless, the previously reported X-ray structure of [4Fe-4S] *Bt*Fd was at 0.92 Å resolution, and all of the hydrogen bonds were invisible. This demonstrated that neutron analysis is an indispensable method for determining the location of hydrogen atoms; the experimentally determined distances/angles are shown in Fig. 2E. Incidentally, depending on the bonding mode in which the S atom is bonded, if the S atom is a hydrogen bond acceptor, the -N-H…S hydrogen bond distance is estimated to be approximately 3.0 Å. In a special case known as a resonance-assisted H-bond, a -N-H…S distance of 2.44 Å has also been reported, in which case a strong hydrogen bond is formed(13). The average distance between the oxygen and the sulfur is reported to be 3.32 Å, and the distance of the nitrogen and sulfur is 10% longer compared to that of the oxygen and sulfur(13); the distance between the nitrogen and sulfur is estimated to be approximately 3.6–3.7 Å.

Importantly, the structural comparison showed that the neutron structure in this study was able to accurately revise the hydrogen network around the [4Fe-4S] cluster (Fig. 2F). The -OH group on the sidechain of Thr63 was located an appropriate distance from the [4Fe-4S] cluster in order to form a hydrogen bond. This group was thought to be hydrogen-bonded in previous X-ray study(12) as the hydrogen atom in this position was not visible, but the distance as well as the angle between the donor and acceptor atoms indicated the possibility of the formation of a sulfur-containing hydrogen bond(14). In the neutron crystal structure, Thr63-Oγ1 and the [4Fe-4S]-S4 atoms were also within the hydrogen-bonding distance (3.5 Å), however, the hydrogen atom identified pointed in a different direction from the iron-sulfur cluster, indicating that it did not form a hydrogen-bonding interaction with the iron-sulfur cluster (Fig. 2F). Furthermore, the neutron-scattering length density clearly revealed hydrogen bonding to another residue; the sidechain of Thr63-Oγ1H and the main chain -CO of Thr10 clearly formed a hydrogen bond at a distance of 2.2 Å. In order to consider the redox potential, it is important to experimentally determine the exact hydrogen-bonding network, and, ultimately, a single hydrogen bond can make a critical difference.

### Density functional theory calculations of the low-potential [4Fe-4S] cluster in *Bt*Fd

In this study, we were able to revise the hydrogen-bonding network around the low-potential [4Fe-4S] cluster. We then investigated the effect of hydrogen bonding around the active site by the DFT calculations using the [4Fe-4S] cluster model based on the neutron diffraction structure including some amino acid residues (**CM**), as shown in Fig. 3A. The Cartesian coordinates of the model are summarized in Table S1. At first, the charge/spin state of each iron atom in the [4Fe-4S] cluster was examined using a cluster model without amino acid (**CM_NA_**) as shown in Fig. 3A. In accordance with previous reports(15, 16), the total energies of the 12 possible combinations of charge/spin alignments were examined for the [Fe^III^_2_Fe^II^ ] (oxidized: *Ox*) and [Fe^III^Fe^II^ ] (reduced: *Red*) states as summarized in Table S2. As shown in Fig. 3B, the most stable spin alignments for the *Ox* and *Red* states are confirmed to be consistent with the states previously reported(16), thus we assumed those charge/spin alignments in the following calculations. Next, to estimate the effect of hydrogen bonding, the vertical ionization potential (IP) was estimated by the calculated energy gaps between oxidized [*E*(*Ox*)] and reduced [*E*(*Red*)] states(16, 17). Note that the IP value is obtained for each model by calculating both the *Ox* and *Red* state energies of the model.

**Fig. 3.**
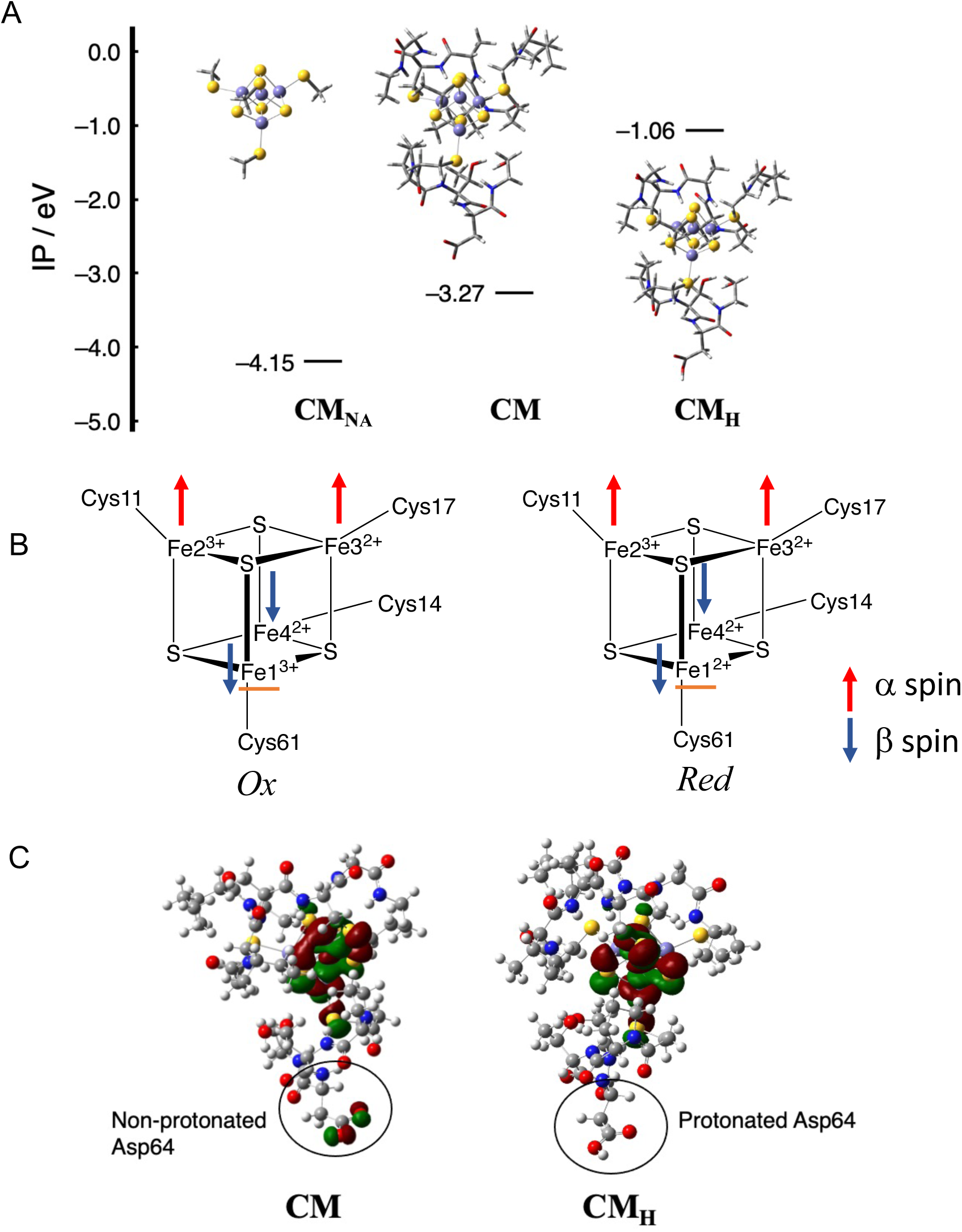
DFT calculation based on the neutron structure of [4Fe-4S] *Bt*Fd. (*A*) Illustration of model structures (CM_N_, CM, and CM_H_) and their calculated *IP* values. (*B*) Assumed charge/spin states of the [4Fe-4S] cluster for Ox and Red states. Fe1 is indicated by the orange underscore and dominantly changes its charge state. (*C*) Distribution of LUMOs of CM and CM_H_. The red and green colors indicate positive and negative phases, respectively.

The calculated IP value of **CM** was −3.27 eV, suggesting the *Ox* state was 3.27 eV more stable than the *Red* state in this structure. In order to consider the effect of the surrounding amino acids, the IP value was also calculated using **CM_NA_**. The calculated IP value was −4.15 eV, which was shifted 0.88 eV to the negative side in comparison with **CM**, suggesting that the surrounding amino acids affect the IP value. In the previous study(18), it was reported that [N–H ⋯ S] hydrogen bonds involved with the cluster stabilize the reduced state and shift the IP of the cluster to the positive side. Our result was, therefore, consistent with the previous study(18). However, a single [N–H ⋯ S] hydrogen bond is suggested to change the IP value approximately 0.3 eV(18), so the 0.88eV shift was much smaller than the value expected from the neutron structure (9 hydrogen bonds) around the active site. To explain this underestimation in the IP value, the lowest unoccupied molecular orbital (LUMO) of the oxidized state, which was the corresponding orbital in this redox reaction, was examined as illustrated in Fig. 3C. The LUMO was mainly localized around the 4Fe-4S cluster; however, unexpectedly appeared at the deprotonated Asp64. This means that the electron density is partially removed not only from the [4Fe-4S] cluster but also from

Asp64 in the oxidation process (*Red* → *Ox*). We note that the carboxy group of Asp64 is not neutralized (–COO^−^) in the neutron structure, but it is easily protonated and neutralized since Asp64 is on the surface of the protein. Therefore, the IP of the cluster was recalculated with the **CM_H_** model as illustrated in Fig. 3A. Then the population of LUMO around Asp64 was removed and fully localized on the [4Fe-4S] cluster as illustrated in Fig. 3C. Interestingly, the calculated IP of **CM_H_** (–1.06 eV) was largely shifted to the positive side in comparison with the deprotonated **CM**.

The difference in the IP value between **CM_H_** and **CM_NA_** (3.09 eV) indicates a contribution of 0.34 eV per one hydrogen bond, which is comparable to the previous report(18). These results suggest that the protonation state of the sidechain of Asp64 plays important roles for the stability of the reduced state, and furthermore in the control of electron transfer. To examine the charge difference between **CM** and **CM_H_** in the simulation, we also constructed a new model including an additional OH^−^ ion around Asp64 (**CM_HOH_**) as illustrated in Fig. S2A. The total charge of the **CM_HOH_** model corresponds to the **CM** model and includes the effect of the hydrogen bond. The calculated IP for the **CM_HOH_** is –2.93 eV, therefore a contribution of a single hydrogen bond is 0.33 eV. The decrease in |IP| value indicates that the relative stability of the *Red* state is suppressed compared with the **CM_H_** but is significantly larger than the **CM**, suggesting the importance of the protonation of Asp64 (Fig. S2B).

To consider the effect of the structural change caused by the redox on the IP, geometrical optimization of the 4Fe-4S core was performed for the **CM** (*Red*) and **CM_H_** (*Red*) models using the same level of theory to the single-point calculations. The optimized Cartesian coordinates are summarized in Table S3. As illustrated in Fig. S2A, the IP values of **CM** and **CM_H_** change from – 3.27 to –2.38 eV (|ΔIP| = 0.89 eV), and from –1.06 to –0.19 eV (|ΔIP| = 0.87 eV), respectively, before and after the geometrical optimization. These results indicate that the IP values are significantly affected by the structural rearrangement; however, the shifts (|ΔIP|) are the same between the **CM** and **CM_H_** models. In other words, the effect of the structural change on the redox in the IP value is important but the significance of the protonation to Asp64 is not changed, regardless of the structural rearrangement.

To consider the environmental effect of the surrounding protein, we also introduced The Polarizable Continuum Model (PCM) using the Integral Equation Formalism variant (IEFPCM) approximation. As summarized in Table S4 in detail, IP values of **CM** and **CM_H_** becomes 1.62 eV and 2.59 eV, respectively, indicating that the environmental effect significantly changes the IP value. On the other hand, a qualitative relationship between the IP values of the **CM** and **CM_H_** models is not changed, although the difference in the IP values between **CM** and **CM_H_** (ΔIP) is decreased from ca. 2.2 to 1.0 eV. From these results, we can conclude that the protein environment is essential for their IP values but does not change the relative difference between **CM** and **CM_H_** (ΔIP) qualitatively.

### The role of Asp64 in *Bt*Fd for the air-oxidation of the low-potential [4Fe-4S] cluster

The DFT calculations highlighted the influence of the Asp64 protonation state on the IP value of the [4Fe-4S] cluster. The IP value is directly reflected in the oxidation rate of the [4Fe-4S] cluster in atmospheric air, since the electron transfer rate to oxygen depends on the IP. In *Bt*Fd, the ultraviolet-visible (UV-Vis) spectra were changed between reduced and oxidized states of the [4Fe-4S] cluster (Fig. 4A); the absorption shoulders around 320 nm and 420 nm were increased by oxidation. Therefore, we prepared reduced *Bt*Fd under anaerobic conditions and measured the spectral changes at 420 nm, associated with air oxidation.

**Fig. 4.**
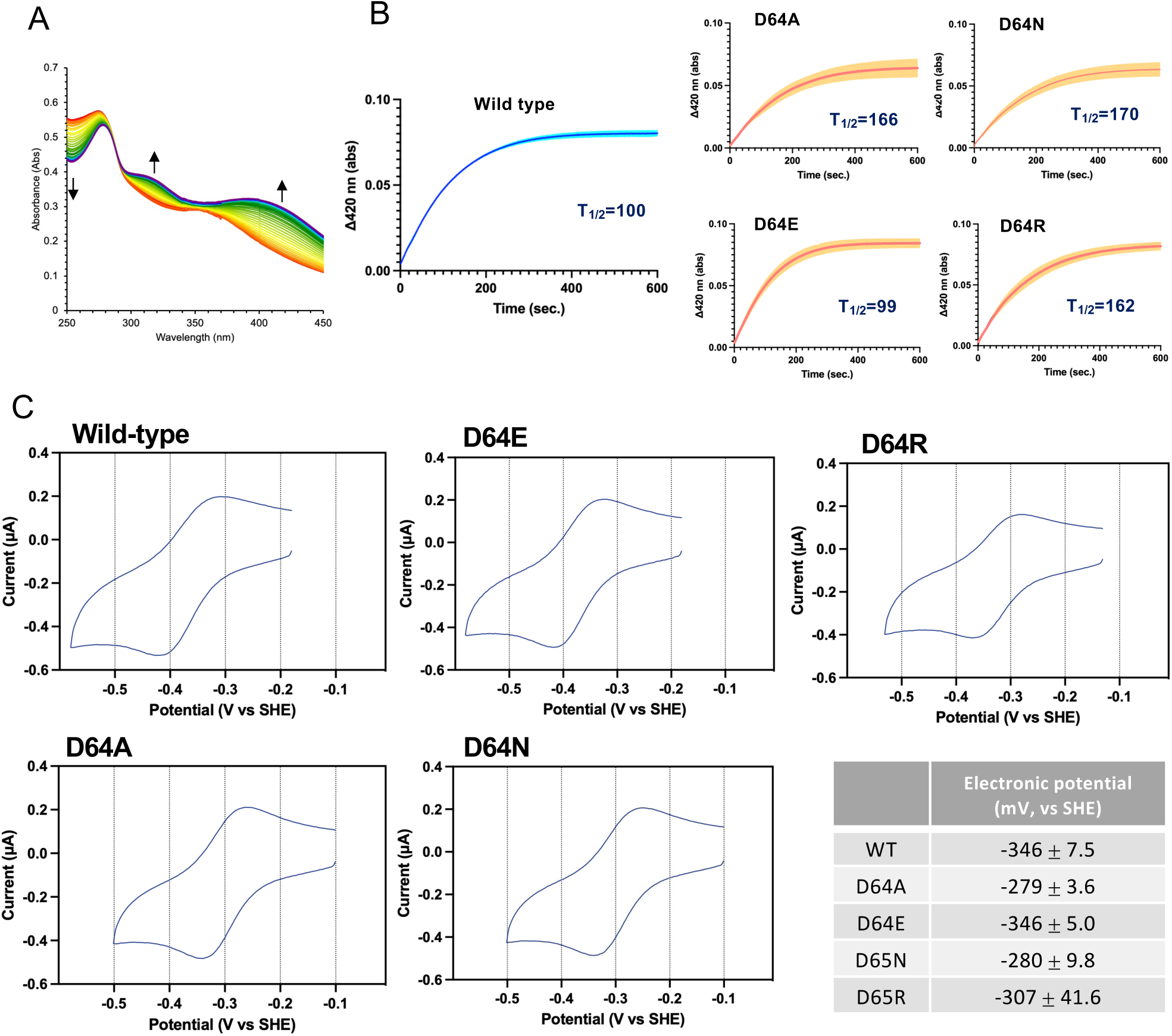
Spectral changes upon [4Fe-4S] cluster oxidation and typical voltammogram of *Bt*Fd and its mutant proteins. (*A*) Time-dependent changing of UV-vis absorption spectrum of wild-type *Bt*Fd by air oxidation. The spectra are colored in rainbow colors from red (reduced BtFd) to purple (oxidated BtFd). The arrows indicate the absorbance increase at 340 nm, 420 nm, and indicate the absorbance decrease at 260 nm. (*B*) The absorption changes of the Asp64 mutated BtFds at 420 nm recorded every second. The standard deviations calculated from at least 3 times the measurements indicated as the line width with the light color. The T_1/2_ value (sec) is indicated in the inset. (*C*) The cyclic voltammograms obtained at the disposable screen-printed carbon electrode from a solution of as isolated 100 μM *Bt*Fd or its mutated proteins in 50 mM Tris-HCl (pH 7.0), 150 mM NaCl and 2.5 mM neomycin, at sweep rate of 50 mV/s. The CV measurement performed in the anerobic chamber (O_2_ < 5 ppm ).

In wild-type *Bt*Fd, the time-dependent change of the UV-Vis spectrum is associated with oxidation of the [4Fe-4S] cluster and displayed typical Michaelis-Menten kinetics (Fig. 4B). Therefore, we applied the global fitting to evaluate the oxidation rate (half time of maximum oxidation (T_1/2_) as for the *K*_m_ in substrate affinity), and in the evaluation, small numbers indicated the fast oxidation reaction. Multiple measurements of the oxidation rates showed a T_1/2_ of 100 sec for wild-type *Bt*Fd. In contrast, that of D64A-mutated *Bt*Fd was 166 sec, approximately 1.7 times slower than wild type. The D64N mutant also had a slow oxidation rate at 170 sec. These results clearly indicate that a neutralizing mutation in Asp64 leads to a slow oxidation rate, and the reduced state of the [4Fe-4S] cluster has greater stability than that of wild type.

*Bt*Fd has 15 aspartate residues including Asp64, all of which are located in the solvent-exposed environment (Fig. 1). We mutated each aspartate to alanine and evaluated changes in oxidation rates; however, only mutations in Asp64 resulted in oxidation rate changes (Fig. 4B and Fig. S3). *Bt*Fd also has 5 glutamate residues, but none of these affected the oxidation rate (Fig. S3).

More interestingly, the D64N neutralizing mutation had the same effect on [4Fe-4S] as the D64A mutation, and the D64R mutation, a positively charged mutant, also slowed the similar oxidation rate (Fig. 4B). In contrast, the D64E mutation, a negatively charged mutation, did not affect the oxidation rate. Namely, the carboxy group in this position exposed to solvent is indispensable for the function of the electron transfer.

### The role of Asp64 in low-potential [4Fe-4S] *Bt*Fd for the redox potential

To further evaluate the effect of the mutation, we determined the redox potential (E^°^, V versus standard hydrogen electrode (SHE), pH 7.0 of each mutant by the cyclic voltammetry method; Fig. 4C). As a result, the redox potential of wild-type *Bt*Fd was estimated to be -346 mV. In contrast, the potentials of the D64A and D64N-mutated *Bt*Fd were –276 mV and –286 mV, respectively. The potential of the D64E mutation was –346 mV, similar to that of the wild type. The neutralizing mutation in the Asp64 position causes the positive shift of 60 ∼ 70 mV in redox potential. The potential of the D64R mutation was estimated to be -307 mV, but its deviation is large (± 41.6 mV).

### The effect of aspartate/glutamate residue in various ferredoxins harboring the low-potential [4Fe-4S] cluster

We performed further experiments to confirm whether the results obtained with *Bt*Fd are the same for various ferredoxins from different organisms. We attempted to overexpress and purify eight ferredoxins as in the phylogenetic tree (Fig. 5A) from Eukaryote, Archaea, and Bacteria, in which three of them were successfully obtained from the ferredoxins harboring the [4Fe-4S] or [3Fe-4S] cluster. The [4Fe-4S] ferredoxin from *Staphylococcus haemolyticus,* a close relative to *Bacillus thermoproteolyticus*, has an aspartate residue at the position of 65 (Fig. S4C), which corresponds to Asp64 in *Bt*Fd. The measurement of the air-oxidation rate indicated that the Asp65 mutation critically affected the rate as seen in that of *Bt*Fd (Fig. 5B). In the *Megalodesulfovirio* (*Desulfovibrio*) *gigas* [3Fe-4S] ferredoxin, the Glu53 is the corresponding position to the Asp64 in *Bt*Fd (Fig. S4C). The air-oxidation rate is quite slow and its rate is further slowed by introducing the E53A mutation (Fig. 5B). Surprisingly, even in the archaeal *Geoglobus acetivorans* [4Fe-4S] ferredoxin, which is phylogenetically quite distant from *Bt*Fd, the E56A mutation affected the oxidation rate (Fig. 5B).

**Fig. 5.**
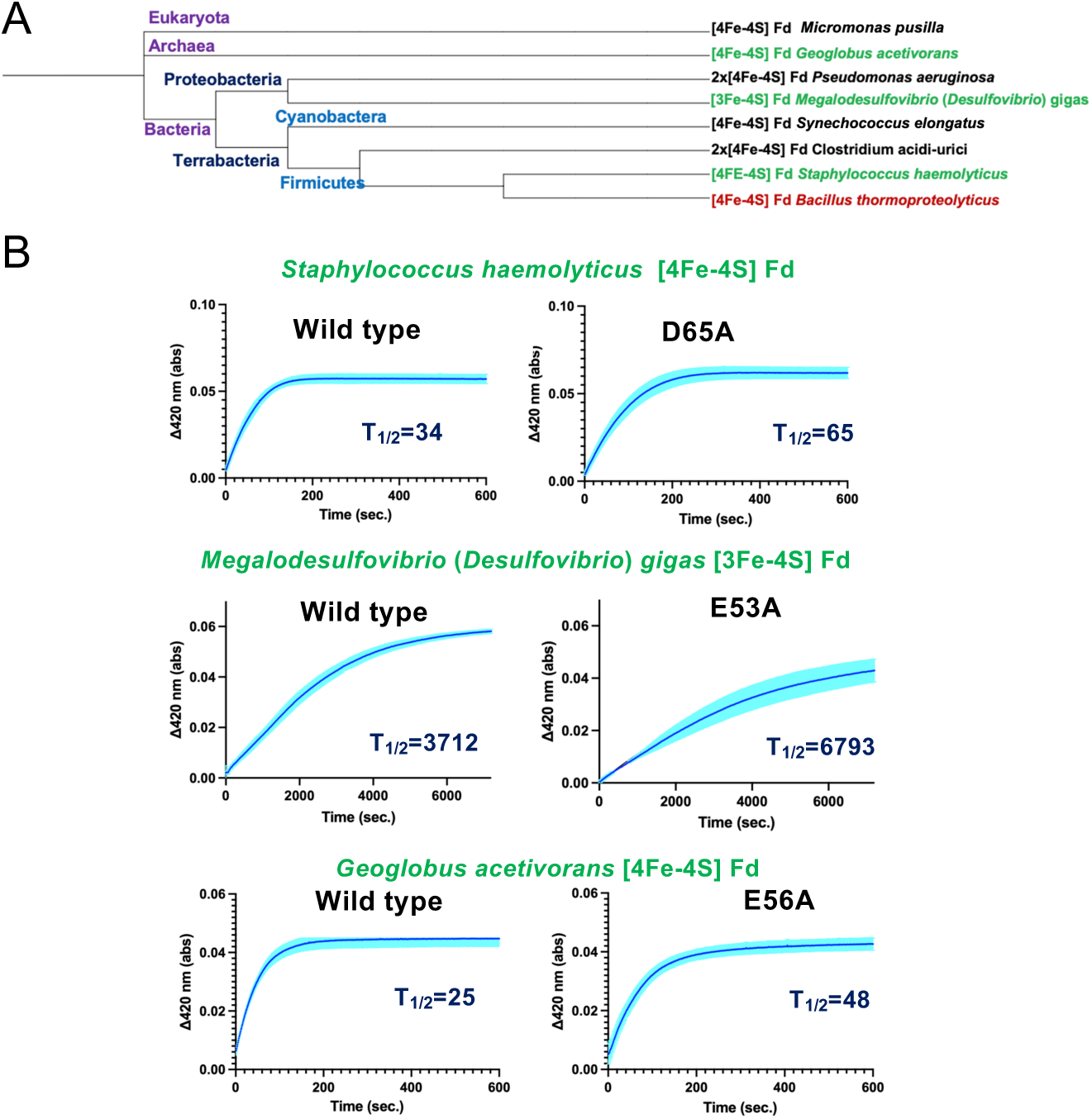
The effect of aspartate/glutamate residue in various ferredoxins harboring the low-potential [4Fe-4S] cluster. (A) Pyrogenetic tree of ferredoxin harboring the [4Fe-4S], [4Fe-4S] x 2 and [3Fe-4S] cluster from various organism. (*B*) The air-oxidation absorption changes at 420 nm of various ferredoxin and its Asp/Glu mutated ferredoxin are also indicated.

## Discussion

Redox potentials are major contributors to controlling electron transfer rates and thus regulating electron transfer processes in biological reactions. Elucidating how proteins control the redox potentials of Fe–S clusters is a longstanding fundamental question. Possible controlling factors that have been proposed as being important include the solvent exposure of the cluster, specific hydrogen-bonding networks, especially NH…S bonds, and the proximity and orientation of the protein backbone and sidechain dipoles.(7, 19–22) Thus far, several mutagenesis studies have demonstrated that the redox potential of ferredoxin can be changed by introducing artificial mutations into residues near Fe-S clusters. For instance, in [4Fe-4S] ferredoxin from *Azotobacter vinelandii* (*Av*Fd I), mutations in positions Phe2 or Phe25 showed a large shift (100–200 mV versus SHE) in reduction potentials(23). The series of mutations based on the sequence/structural comparison among *Av*Fd I and *Peptostreptococcus asaccharolyticus* ferredoxin (*Pa*Fd) and related ferredoxins showed that three mutations, V19E, P47S, and L44S in *Av*Fd I also shift to higher reduction potentials (130 mV increasing in total)(19). These artificial mutation studies clearly showed the significance of the environment of [4Fe-4S] for redox potentials; however, their intrinsic control residues have remained ambiguous.

In this study, we showed that Asp64 in the *Bt*Fd affected the behaviors of the [4Fe-4S] cluster including the redox potential (Fig. 4B and C). During the course of the purification, all of the mutated *Bt*Fd eluted at the same conductivity from the high-resolution ion-exchange column, that is, the aspartate mutation in various positions did not disturb the net charge of this protein. Also, the respective UV-Vis spectra of the reduced and oxidized forms of *Bt*Fd mutants were the same as that of the wild type (Fig. S4), demonstrating that the environment of the [4Fe-4S] cluster was not altered by these mutations. These results indicated that the acidic residue mutations in other positions did not significantly affect the oxidation rate (Fig. 4B). The distances between the Asp/Glu and [4Fe-4S] cluster are listed in Table S5; the sidechain of Asp64 is located at 10.1 Å, and the Asp7 (10.5 Å), Asp26 (12.0 Å), Asp28 (11.3 Å), Asp30 (13.7 Å), Asp53 (13.7 Å), Glu9 (13.7 Å), and Glu59 (12.8 Å) were located at similar distances. Taken together, only Asp64 affects the oxidation rate of the [4Fe-4S] cluster, and the protonation state of Asp64 directly contributes to the control of electron potential of the [4Fe-4S] cluster.

The mutation experiments showed that the Asp64 residue in *Bt*Fd was exchangeable with a glutamate residue, which means that the carboxy group in this position is indispensable. The sequence alignment of the low-potential [4Fe-4S] ferredoxin demonstrated that negatively charged residues, aspartate or glutamate, are highly conserved at this position among low-potential [4Fe-4S] ferredoxins in bacteria (Fig. S6). Interestingly, the significance of the highly conserved aspartate or glutamate was demonstrated for the ferredoxins from different organisms even in organisms distantly related in the evolutionary phylogenetic tree (Fig. 5A and B). That is, the effect of the D64A mutation in the [4Fe-4S] *Bt*Fd in gram-positive bacteria is also observed in the E56A mutation and in archaeal [4Fe-4S] ferredoxin from *Geoglobus acetivorancs*. Furthermore, the same results are obtained from the [4Fe-3S] ferredoxin from *Megalodesulfovibrio gigas,* a gram-negative bacterium. Thus, the control mechanism of the redox-state stability found in *Bt*Fd may be a common mechanism, at least in low-potential [4Fe-4S] and [3Fe-4S] ferredoxins.

In general, the p*K*a value of the aspartic acid sidechain is a pH of approximately 3.9, meaning that under physiological conditions, the sidechain of Asp64 is constantly in a deprotonated state. We have attempted to determine the p*K*a value of the Asp64 in *Bt*Fd by monitoring the chemical shifts of the Asp64 sidechain ^13^CO(γ) using nuclear magnetic resonance to determine the pH titration of Asp64. Under the reduced conditions we could not determine the p*K*a value of the carboxylate hampering the paramagnetic property of the reduced [4Fe-4S] cluster. Instead, under the oxidative conditions (normal ambient air conditions), the p*K*a of Asp64 was successfully estimated to be approximately 4.9 (Fig. S7A and B, SI method and text). This indicated that the p*K*a of Asp64 shifted to neutral. To further estimate the p*K*a under reduced conditions, we applied the air-oxidation measurement; the oxidation rate of the wild-type *Bt*Fd changed dramatically between pH 6.0 to 7.0 (Fig. S7C). In contrast the D64A *Bt*Fd did not alter the rate according to the pH. These results indicate that the p*K*a value of Asp64 is thought to be between a pH of 6.0 to 7.0. The [4Fe-4S] cluster in the *Bt*Fd was stable enough to conduct these experiments under acidic to basic conditions (pH 4.5–9.0) (Fig. S8). Thus, the protonation/deprotonation of Asp64 likely occurs even in physiological conditions.

The structural basis of the interactions between ferredoxin and its partner proteins has been previously reported(24–26). In the reported structures, ferredoxins bound to their partner proteins such as ferredoxin:NADP^+^ oxidoreductase or photosystem I were required to directly face the Fe-S cluster region to effectively accept electrons from their partner proteins. These complex structures are derived from photosynthetic organisms, and ferredoxins harboring a [2Fe-2S] cluster are called plant-type ferredoxins. Unfortunately, there is no structural report about the characteristics of the interaction between bacterial [4Fe-4S] ferredoxins and their partner proteins, although the major interaction region of ferredoxin must be in close contact with the Fe-S cluster. In bacterial low-potential [4Fe-4S] ferredoxins, if an aspartate residue neighboring the [4Fe-4S] cluster, corresponding to Asp64 in *Bt*Fd, was protonated by complex formation, electrons received from the bacterial ferredoxin:NADP^+^ oxidoreductase complex are able to remain stable. According to the Frank-Condon principle, known as the ground rule of the electron transfer, the electronic transitions are relatively instantaneous compared with the time scale of nuclear motions. As well, in *Bt*Fd, the distance between the [4Fe-4S] cluster and the carboxy sidechain of Asp64 was close enough to enable the electron transfer as the corresponding distance is approximately 10 Å (Table S5). Therefore, the protonation state of the aspartate residue neighboring the [4Fe-4S]/[3Fe-4S] cluster is a reasonable switch to control the stability of the reduced state in the [4Fe-4S]/[3Fe-4S] cluster. We propose a control mechanism of the redox potential in low-potential [4Fe-4S]/[3Fe-4S] ferredoxins as shown in Fig. 6. This protonation-driven switch is likely reasonable to prevent the backflow of accepted electrons.

**Fig. 6.**
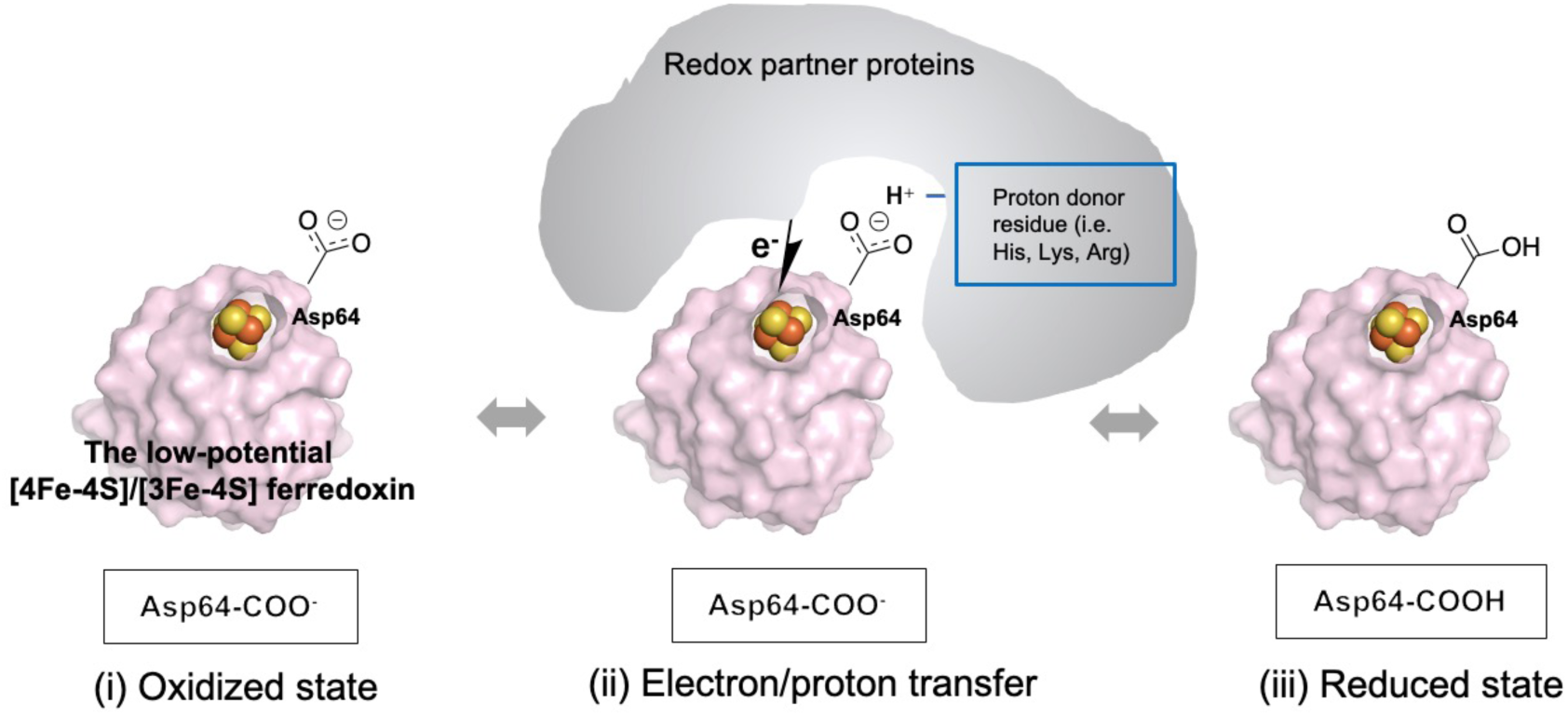
Proposed putative mechanism of the low-potential ferredoxin. (i) The deprotonation state of the aspartate residue (Asp-COO^-^) neighboring the oxidated [4Fe-4S]/[3Fe-4S] cluster is the receivable state of an electron. (ii) When the electron-donor protein forms a complex with ferredoxin, the electrons move to the [4Fe-4S]/[3Fe-4S] cluster instantaneously. (iii) Next, the aspartate residue accepts the proton for neutralization (Asp-COOH), and the reduced state of [4Fe-4S]/[3Fe-4S] clusters is stabilized.

To our knowledge, this is the first proposed mechanism of the intrinsic control factor of redox potentials of the [4Fe-4S]/[3Fe-4S] cluster in low-potential ferredoxins. We hope that the determination of complex structures with electron donors and acceptors involving ferredoxin will elucidate more detailed control mechanisms for electron transfer by low-potential [4Fe-4S]/[3Fe-4S] ferredoxins.

## Materials and Methods

### Expression and purification of the low-potential ferredoxin of [4Fe-4S] *Bt*Fd

Expression and purification of [4Fe-4S] *Bt*Fd were conducted as previously reported(12, 27), with some modifications. The plasmid pET-21a(+)-*btfd* gene was co-expressed with the *isc* operon (pRK-ISC)(28) in the *E. coli* strain C41(DE3) to produce holo-*Bt*Fd in high yield. The bacteria were cultivated in 2.5 liters of Terrific broth containing 100 µg/mL ampicillin, 5 µg/mL tetracycline, and 0.1 mg/mL ferric ammonium citrate. Expression was induced with 0.5 mM IPTG and the cells were further grown for 20 h at 28°C. The cells were disrupted by sonication, and the suspension was centrifuged at 12,000 *g* for 30 min at 4°C. The colored supernatant was subjected to ammonium sulfate fractionation at 60% saturation. After centrifugation, the supernatant fraction was applied to HiPrep Butyl FF 16/10 (Cytiva, Marlborough, MA, USA) and eluted with a linear gradient of ammonium sulfate (5–25%) in 50 mM Tris-HCl, pH 7.8. The grayish brown fractions were dialyzed against a solution containing 50 mM Tris-HCl, pH 7.8, applied to an anion exchange column (Mono Q HR 5/5, Cytiva) chromatography using the NGC Quest 10 system (Bio-Rad Laboratories, Inc. Hercules, CA).

### Expression and purification of the low-potential ferredoxin of [4Fe-4S], [4Fe-4S]x2, and [3Fe-4S] ferredoxin from various organisms

Expressions and purifications of [4Fe-4S] ferredoxins were conducted using the same procedures as that of *Bt*Fd. Each ferredoxin gene was constructed using a synthetic oligonucleotide (Eurofins Japan, Tokyo) based on UniProt database sequence optimized for the *E. coli* codon usage.

### Crystallization of [4Fe-4Fe] BtFd for neutron crystallography

The crystallization procedure and conditions are as follows and were modified from the method reported previously to grow large crystals for neutron crystallography(12). Purified [4Fe-4S] BtFd was concentrated by centrifugation using an Amicon Ultra centrifugal filter unit 3K (Merck Millipore) to 30–40 mg/mL. The crystal for the neutron diffraction experiment was obtained with the dialysis method at 4°C using a 50 μL dialysis button (Hampton Research, Journey Aliso Viejo, CA, USA) and the excess amount of the reservoir solution containing 200 mM NaCl, 1.4 M ammonium sulfate, and 50 mM MES (pH 5.9). The crystal appeared within one month and was grown to approximately 2.5 mm × 1.8 mm × 0.6 mm in a month.

### Neutron diffraction experiment

The crystal was soaked to 50 μL of 50 mM MES buffer (pD 6.3) containing 200 mM NaCl and 1.4 M ammonium-d8 sulfate (D8, 98%; Cambridge Isotope Laboratories Inc.) prior to the neutron diffraction experiment. The soaking solution was exchanged three times in three weeks. The crystal was mounted in a quartz glass capillary with 3.5 mm ϕ and 0.01 mm thickness (Hilgenberg). Inside the capillary, a small amount of deuterated reservoir solution was contained to avoid drying up the crystal, and the capillary was sealed with epoxy resin. Time-of-flight (TOF) neutron diffraction data were collected at BL03 iBIX (29) in a Japan Proton Accelerator Research Complex (J-PARC, Tokai, Japan) at room temperature. Thirty-two wavelength-shifting fiber-based scintillator neutron detectors with a surface area of 133 mm × 133 mm were used to collect the data. A total of 56 data sets were collected using a wavelength of 1.5–4.5 Å with a detector distance of 490 mm. Exposure time for each data set was 6 h at 500 kW of the J-PARC accelerator power. The TOF neutron data were indexed, integrated, scaled, and processed with STARGazer(30). The neutron diffraction data statistics are listed in Table 1.

### X-ray diffraction experiment

X-ray diffraction data from the same crystal used for neutron crystallography were collected using an ADSC Quantum270 CCD detector at NW12A in Photon Factory Advanced Ring (PF-AR; Tsukuba, Japan) under room temperature. The wavelength of the synchrotron radiation, transmittance, and slit size were 1.000 Å, 1%, and 0.20 mm (horizontal) × 0.05 mm (vertical), respectively.^31^ The sample-to-detector distance, oscillation range and exposure time were 75 mm, 0.5° and 0.5 sec, respectively. In total, 720 images were collected. Data were integrated, merged, and processed with HKL-2000(31). The X-ray diffraction data collection and statistics are listed in Table 1.

### Structure refinement

The structure was determined by molecular replacement method using the previously determined X-ray structure of [4Fe-4S] *Bt*Fd (PDB ID: 1IQZ) from which the waters and [4Fe-4S] cluster were removed as the initial model. The first refinement was carried out using the rigid-body refinement program RefMac5(32) from 8.0 to 3.0 Å resolution range using only X-ray intensity data. The *R* factor was dramatically dropped by rigid-body refinement. Then, the joint refinement was carried out with both the neutron and X-ray diffraction data using PHENIX(33). The 1.6 Å resolution neutron data and the 1.45 Å resolution X-ray data were used for the structure refinement. Five percent of the data were selected by PHENIX for cross-validation. The neutron scattering length density and electron density maps were visualized and manual model refinement was carried out using COOT(34). Protonation states of amino acid residues and orientation of water molecules were manually adjusted by watching both the neutron scattering length density and electron density calculated before including the hydrogens (H)/deuteriums (D) on COOT; only the neutron scattering length density appeared for the positions of the H (negative density)/D (positive density) atoms. The temperature factors for all atoms and occupancies for H/D atoms and residues having dual conformations were also refined. The refinement statistics are also listed in Table 1. All structure illustrations in figures except for Fig. 3 were prepared with Pymol(35).

### DFT calculations

All calculations were performed by using the spin-unrestricted (broken-symmetry) hybrid DFT method with the B3LYP functional set. As the basis set, 6-31G* and 6-31+G* were used for [Fe, C, N, O, H] and [S] atoms, respectively, for the IP calculations. Only the position of a hydrogen atom bound to a carboxy group of Asp64 in CM_H_ was optimized using the 6-31G [Fe, C, H], 6-31G* [N, O], and 6-31+G* [S] basis sets while the other atoms remained fixed. All of these calculations were carried out in gas phase conditions using the Gaussian 09 program package(36).

The IP was defined as the vertical ionization potential values of the reduced state calculated by IP =*E*(*Ox*)–*E*(*Red*), where *E*(*Ox*) and *E*(*Red*) were calculated as total energies of the oxidized (*Ox*) and reduced (*Red*) states by the DFT method(16, 17).

Cartesian coordinates of calculated models are summarized in Table S1. From the formal charges of Fe, S and other ligands, we assumed that the total charges of the **CM_NA_** and **CM** models are –3 and –4 for oxidized (*Ox*) and reduced (*Red*) states, respectively, whilst total charges are –2 and –3 for the *Ox* and *Red* models in the case of the **CM_H_** model. The **CM_HOH_** model is constructed by placing the OH^−^ ion close to Asp64 in the initial structure, followed by a geometric optimization of only the OH^−^ fragment for both the reduced and oxidized states. To examine the effect of the OH^−^ in detail, the OH^−^ fragment is moved along the O–H axis of the OH^−^ fragment in 0.5Å step. The spin states of *Ox* and *Red* states are the open-shell singlet and doublet states, respectively. The calculated Mulliken charges and spin densities of [4Fe-4S] cores are added in Table S2. As mentioned in the main text, the most stable spin/charge alignment of Fe ions were examined among 12 possible spin/charge combinations using **CM** model, and all other following calculations were only assumed to be these ground states (see Table S2A and B). It is reported that the molecular charges sometimes cause the error especially under approximation of the solvation effect by PCM(37), however the simulation is performed in the gas phase, therefore such effect is assumed to be negligible.

### Oxidation rate analysis of [4Fe-4S] *Bt*Fd and its mutated proteins

Site-directed mutagenesis was carried out using a KOD -plus-Mutagenesis Kit (Toyobo), with a pET21a(+)-containing *Bt*Fd gene as a template. Sequence analysis verified that the resultant constructs were free of errors. Expression and purification of mutated [4Fe-4S] *Bt*Fds were performed in the same way as that for wild type.

Prior to the oxidation rate measurement, we performed a complete reduction of [4Fe-4S] *Bt*Fd by using dithionite; 1 mM (final concentration) dithionite was added to 100 μL of [4Fe-4S] *Bt*Fd (6.0 mg/ml) in anaerobic conditions (O_2_ < 5 ppm; COY). After 1 h of incubation at 4 °C, the dithionite was removed by size-exclusion chromatography (HiTrap Desalting 5 ml column; Cytiva) equipped with the AKTA START system under anaerobic conditions. Complete removal of the dithionite was able to be confirmed with the chromatogram, which indicated the two separated peaks derived from the reduced [4Fe-4S] *Bt*Fd and dithionite. The fraction containing reduced [4Fe-4S] *Bt*Fd (1 mL) was moved into quartz cells with the screwcap under anerobic conditions, and taken out of the anaerobic chamber. Then, the UV-Vis spectroscopic measurement (V730BIO, JASCO) was started immediately after the screwcap was opened. The spectral changes at 420 nm were measured every second for 10 mins, and the measurements were performed three times, at least for the global fitting. The temperature of the cell chamber in the spectrometer was strictly controlled at 28 °C by the Peltier water-cooled cell-holder and the cell solution was continuously mixed by a micro-stirrer. The global fittings of the Michaelis-Menten curve were performed using the kinetics curve fitting mode in the GraphPad Prism 9 (GraphPad Software, San Diego, CA USA, www.graphpad.com).

### The electrochemical measurement of [4Fe-4S] *Bt*Fd and its mutated proteins

To exchange the buffer, the purified *Bt*Fd and its mutated proteins were precipitated by ammonium sulfate and the precipitate was suspended in the buffer at various pHs; the sample solutions contained 100 μM *Bt*Fd, 625 mM NaCl, 2.5 mM neomycin, and 50 mM Tris-HCl at a pH of 7.8 or 50 mM MES-Tris mixed buffer at a pH of 6.0–9.0. The cyclic voltammetry was performed using the μStat-I 400 bipotentiostat (Metrohm AG, Switzerland) controlled by Drop-View 8400 software. A disposable screen-printed carbon electrode (DRP-110, Metrohm AG) was formed by a carbon ink and acted as a working electrode (4 mm diameter). Voltammetric measurements used Ag as a pseudo-reference electrode and carbon ink as a counter electrode. Five cyclic scans were recorded between −0.80 and –0.40 V at a 0.05 V s^−1^ sweep rate. The CV measurements were performed anaerobically by setting the devices, including the potentiostat and the disposable electrode, in an anaerobic chamber (O_2_ < 5 ppm; COY).

### Data availability

The atomic coordinates and structure factors for the neutron structure of the *Bt*Fd at room temperature have been deposited in the PDB under accession code 7YL8.

## Acknowledgements

We would like to express our deepest appreciation to Research Fellows Ms. Makiko Ishihara and Dr. Kenji Ite for their many suggestions and advice. We thank Dr. Yu Hirano of the Institute for Quantum Life Science for the helpful advice on the neutron crystal structure analysis. We also thank members of iBIX at J-PARC of Japan for the neutron diffraction data collection. The neutron diffraction experiments, including the test experiments, were conducted under Proposals 2016PX0003, 2017PX0004, 2019PX3020, and 2020PX3006 (Ibaraki Prefecture Project) in J-PARC. We thank members of the Structural Biology beamlines of PF and SPring-8 for the X-ray diffraction data collection, including preliminary experiments. The X-ray diffraction experiments were conducted under Proposals 2017G561 and 2019G512 in PF and 2022A2724 in SPring-8. This work was partly supported by JSPS KAKENHI numbers 23K17981 (to K.W.), 21H5260 (to K.W.), 20H03196 (to K.W.), 16K07261 (to M.U.), 21K05016 (to S.M.), 22H02562 (to M.M.), 21H01951(to Y.K), 22H02050 (to Y.K.), Ibaraki Prefecture’s Sendokenkyu from 2016 to 2021 (to M.U.), The Takeda Science Foundation (to K.W.) and the Japan Foundation for Applied Enzymology (to K.W.). This research was partially supported by Research Support Project for Life Science and Drug Discovery (Basis for Supporting Innovative Drug Discovery and Life Science Research (BINDS)) from AMED and the Frontier Science Research Center, University of Miyazaki. We also thank T. Yokoyama, T. Kawaguchi, A. Yoshida and Y. Kawagoe of the University of Miyazaki for technical assistance.

## Author contributions

K.F., K.W and M.U. initiated and directed this study together with Y.K., M.M. K.W, K.K Y.I. and M.M prepared proteins. K.K., Y.I., K.K, N.Y and M.U. performed neutron and/or X-ray structural biology experiments. I.E., T.K. and Y.K. performed computational studies of the DFT calculations. M.M., K.I, M.S, K.F. and K.W performed the biological experiments. All authors participate in the data analysis and manuscript preparation.

