## Supplemental_Figs for "Protonation/deprotonation-driven switch for the redox stability of low-potential [4Fe-4S] ferredoxin"

**This PDF file includes:**

Supporting text
Figures S1 to S9
Tables S1 to S5
SI References

**Supplementary Text**

**pKa estimation of Asp64 in BtFd by NMR spectroscopy**

The stable isotopically labeled BtFd proteins were expressed in a 2 L M9 minimal medium containing 0.5 g/L $^{15}NH_4Cl$, with 4 g/L of unlabeled glucose, 1 g/L of $^{13}C_6$-glucose for $^{15}N$, or $^{13}C/^{15}N$ labeled as sole nitrogen and carbon sources, respectively. After induction, the cultures were incubated overnight at 25 °C, and the cells were harvested by centrifugation. The purification was performed in the same manner to the crystallization. The NMR sample for standard heteronuclear experiments was dissolved in 50 mM K-phosphate buffer (pH 6.9) containing 50 mM KCl and 7% $D_2O$. The final concentrations of BtFd in the samples were 0.9–1.5 mM in a final volume of 300 mL. Microcell NMR tubes (Shigemi) were used. All NMR spectra were processed with the software package NMRPipe (*1*) and analyzed using Sparky (*2*). All NMR experiments were performed at 303 K on a Bruker AVANCEIII(HD)600 instrument equipped with a TCI CRYOPROBE. As a first step for analyzing the p*K*a of the aspartate sidechains, we established the resonance assignments by standard heteronuclear multidimensional NMR technique (*3*). The backbone resonance assignments were obtained from the following two- and three-dimensional experiments: 2D $^{1}H$-$^{15}N$ HSQC, 3D HNCO, HN(CA)CO, CBCA(CO)NH, and HNCACB. The sidechain aliphatic $^{1}H$ and $^{13}C$ assignments were obtained from the following three- and four-dimensional experiments: 3D C(CO)NH, 3D H(CCO)NH, 3D HCCH-TOCSY, and 4D HC(CO)NH (*4*). Based on the obtained aliphatic assignments, the $^{13}CO(g)$ resonance assignments of the aspartate sidechains were made by linking the adjacent $^{13}Cb$ chemical shifts. Since the CO signals in the 1D $^{13}C$ experiment were heavily overlapped, we used the 2D CACO experiment (*5*) in which the $^{13}Ca$ offset of indirect dimension was shifted to $^{13}C\beta$ (45 ppm) for monitoring the titrated signals. Namely, we diverted a CACO experiment originally used for correlations of backbone $C\alpha$ and CO to the sidechain CO assignment. To avoid the signal splitting caused by $^{13}C$-$^{13}C$ spin coupling during $^{13}CO$ acquisition, low-bash homonuclear decoupling (*6*) was achieved in the experiments. The pH 3.5 buffer was composed of 50 mM phosphate and 50 mM KCl, the pH 4.0–5.0 buffer was composed of 50 mM acetate and 50 mM KCl, the pH 5.5–6.5 buffer was composed of 50 mM MES and 50 mM KCl, the pH 7.0–8.0 buffer was composed of 50 mM HEPES and 50 mM KCl, and the pH 8.5–9.0 buffer was composed of CHES and 50 mM KCl. These buffers contained 7% $D_2O$ for the lock signal. The p*K*a(s) were estimated from the Hill equation by nonlinear least-square curve fitting using Microsoft Excel implemented SOLVER module (Microsoft).

BtFd was stable enough to conduct the NMR experiments under acidic to basic conditions (pH 3.5–9.0), thus, we were able to estimate the p*K*a of aspartate sidechains by monitoring the chemical shifts of $^{13}CO(\gamma)$ (Fig. S7A). Notably, $^{13}CO(\gamma)$ resonance of Asp64 is isolated from the others, and showed remarkable chemical shift changes upon pH titration (Fig. S7, A and B). The sigmoidal curve was obtained from the Asp64 signals and the p*K*a was estimated to be 4.9 (Fig. S7B). The p*K*a value of Asp74 was also able to be estimated to be 5.0, although its chemical shift change was smaller than that of Asp64. The p*K*a values of the other residues were not able to be evaluated, since they showed no sigmoidal curves in the pH range of 3.5–9.0, or their signals were not reliably assigned. Interestingly, the p*K*a value of Asp74 was also shifted to a natural pH (Fig. S7B). This Asp74 is located away from the [4Fe-4S] cluster at a distance of 22.4 Å (Table S5), and the distance between Asp64 and Asp74 is approximately 10.1 Å. The function of Asp74 is unclear at this stage.

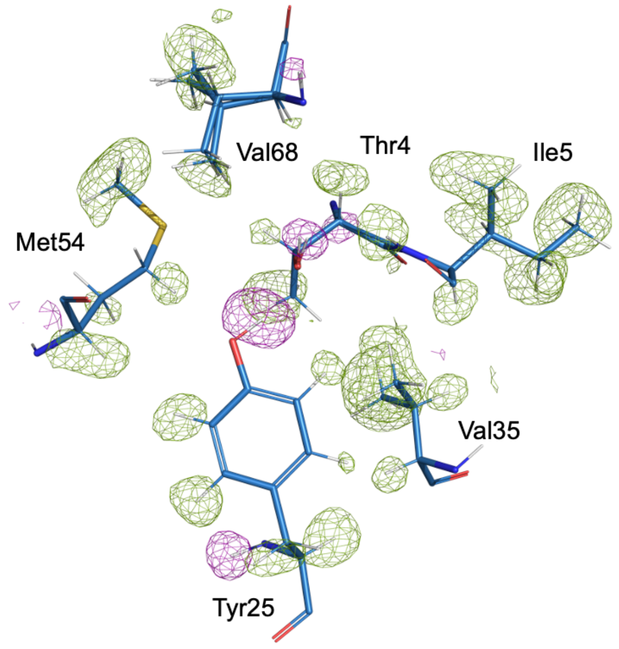


**Fig. S1. Neutron-scattering length density maps of [4Fe-4S] ferredoxins.**
The $F_o$–$F_c$ neutron-scattering length density omits maps for the hydrogen/deuterium at 2.0 σ contour levels. The pink and green cages represent the +($F_o$–$F_c$) and –($F_o$–$F_c$) map, respectively, and the final model of *Bt*Fd is superposed on the maps. The maps clearly indicate that the densities derived from the deuterium and hydrogen atoms are clearly visible in the neutron crystallography.

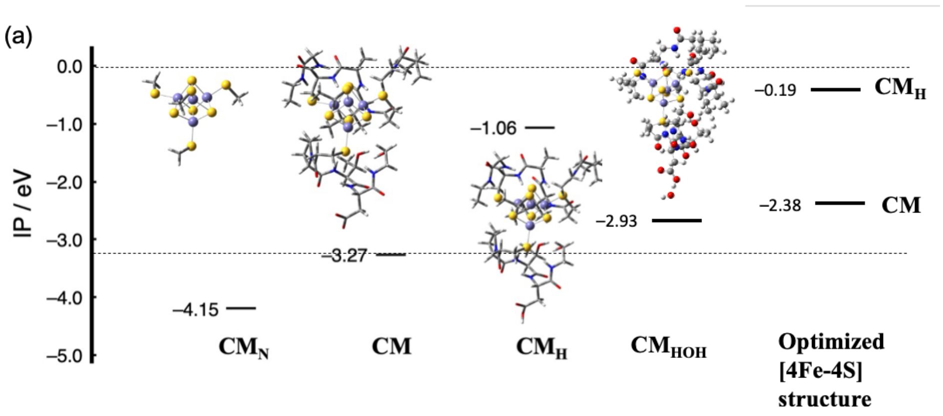


**Fig. S2. DFT calculation based on the neutron structure of [4Fe-4S] BtFd Calculated IP values of each model.**

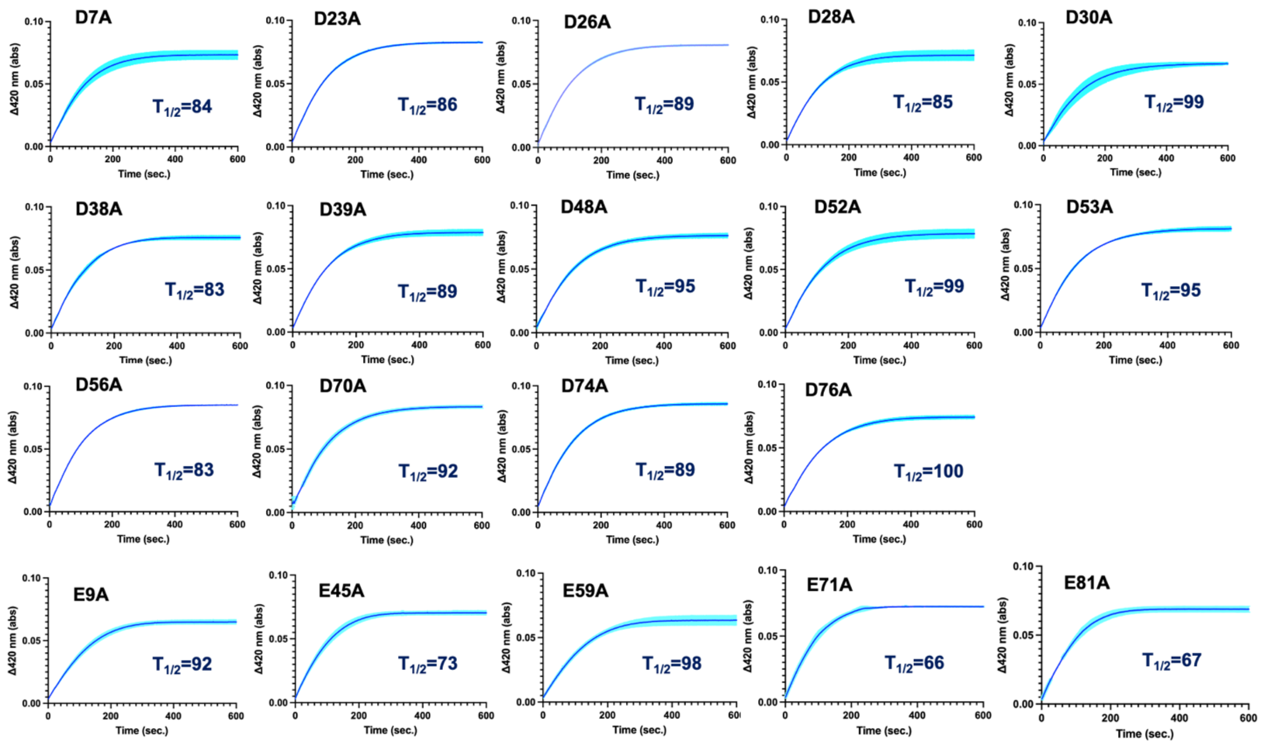


**Fig. S3. The absorption changes of various mutated *Bt*Fds at 420 nm recorded every second.** The standard deviations calculated from at least 3 times the measurements indicated as the line width with the light color. The $T_{1/2}$ value (sec) is indicated in the inset.

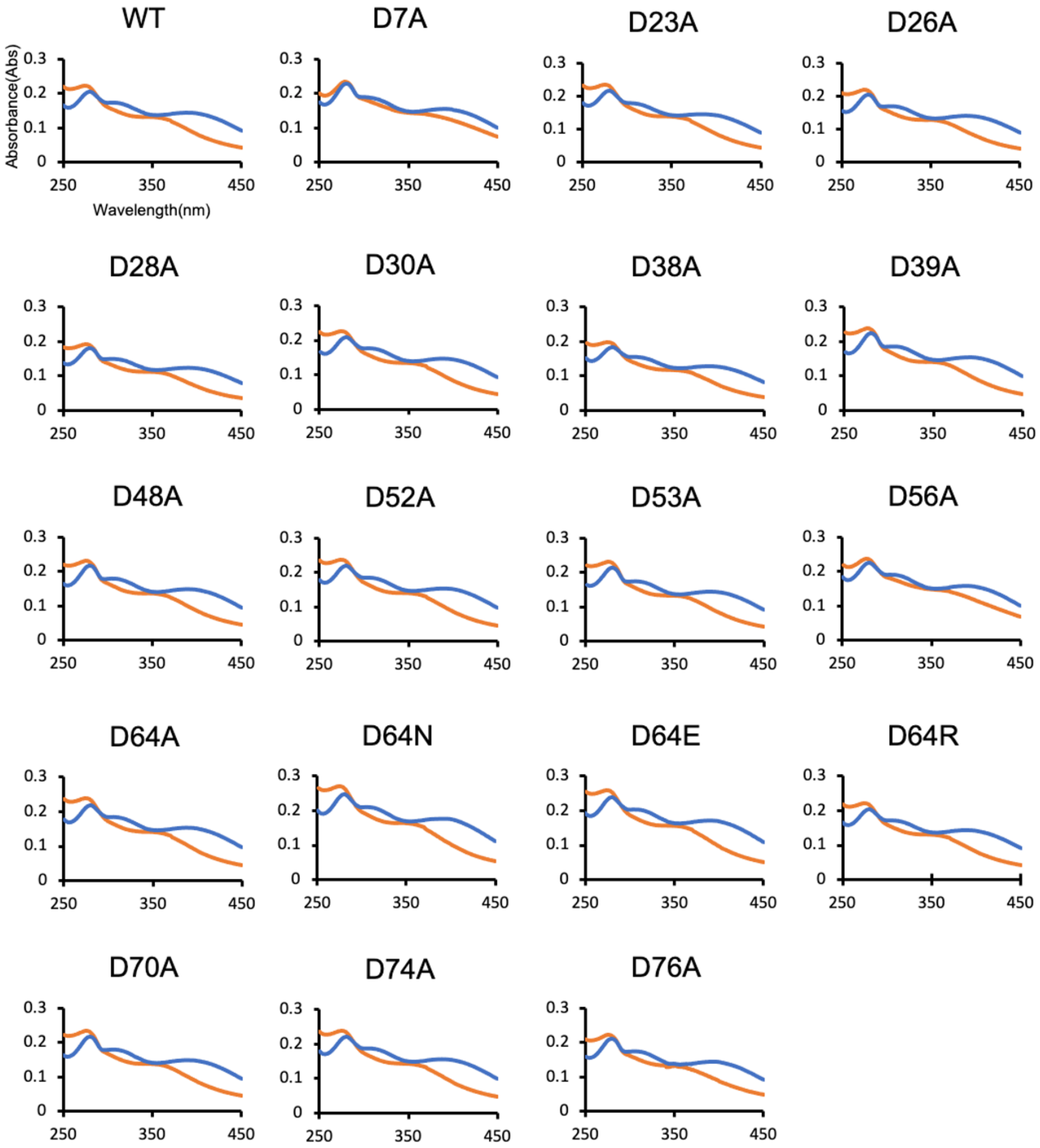


**Fig. S4. UV-Vis absorption spectroscopy of *Bt*Fd and its mutant proteins.**
The spectra of reduced *Bt*Fds are indicated by the orange line and those of completely air-oxidated *Bt*Fds are indicated by the blue line. Wild-type *Bt*Fd and its mutated proteins showed very similar spectra.

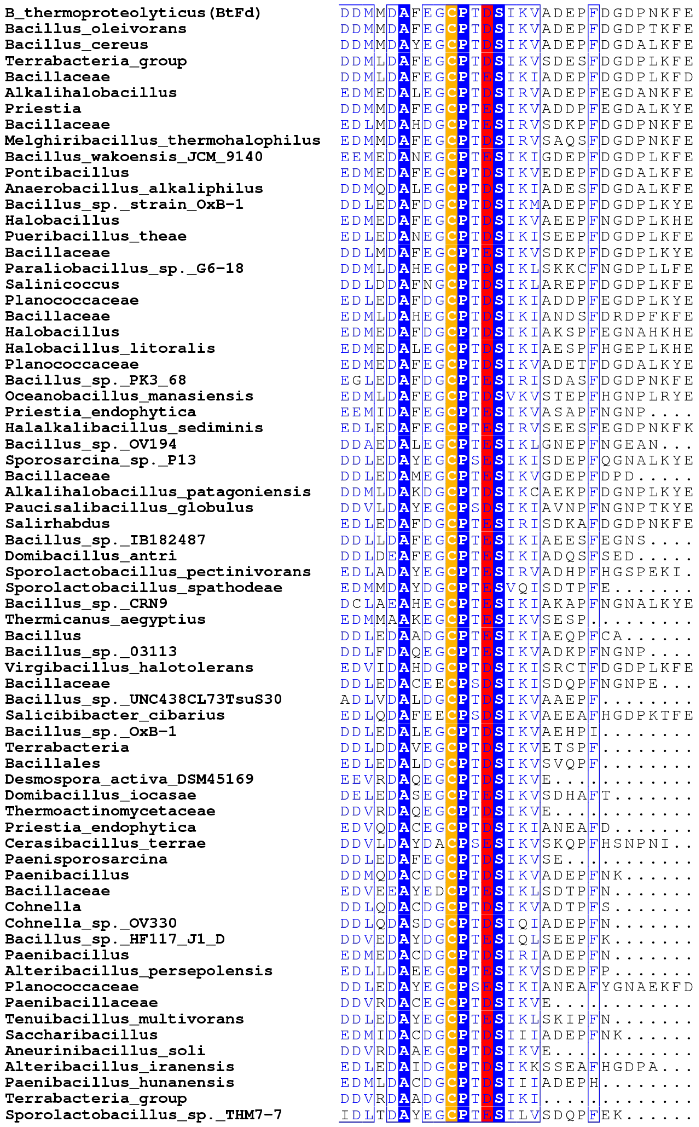


**Fig. S5. Conserved amino acid residues of [4Fe-4S] ferredoxins from several species.** Cysteine residues for the ligand for the Fe-S cluster are completely conserved (orange-highlighted) and several residues around the cysteines are also highly conserved (blue-highlighted). The corresponding residues of Asp64 in BtFd are conserved in negatively charged residues (Asp or Glu: red-highlighted). This figure is prepared using ESpript ver3.0 server (https://espript.ibcp.fr/ESPript/ESPript/index.php).

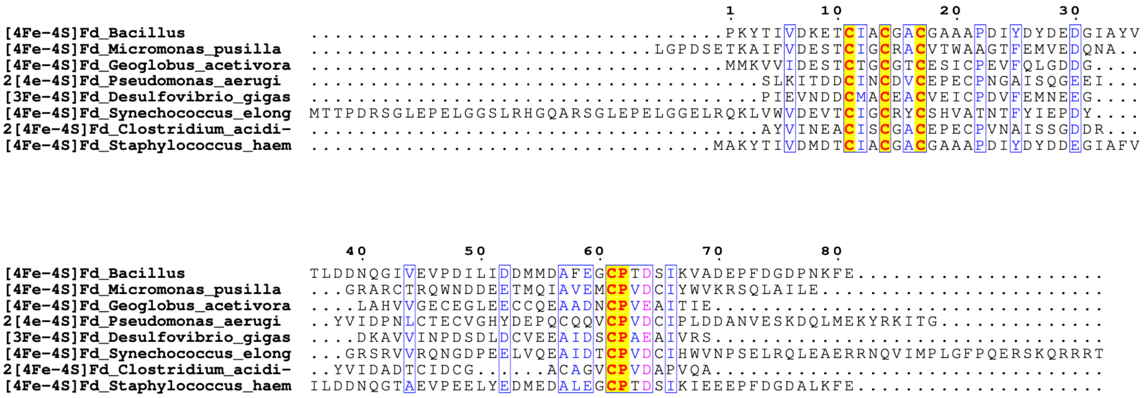


**Fig. S6. Sequence alignment of [4Fe-4S], [4Fe-4S]x2 and [3Fe-4S] ferredoxins from various organisms.**

Cysteine residues for the ligand for the Fe-S cluster are completely conserved (yellow-highlighted) and several residues around the cysteines are also highly conserved (blue-highlighted). The corresponding residues of Asp64 in BtFd are conserved in negatively charged residues (Asp or Glu: pink font). This figure is prepared using the Consurf server (https://consurf.tau.ac.il/index_proteins.php) and ESpript ver3.0 server (https://espript.ibcp.fr/ESPript/ESPript/index.php).

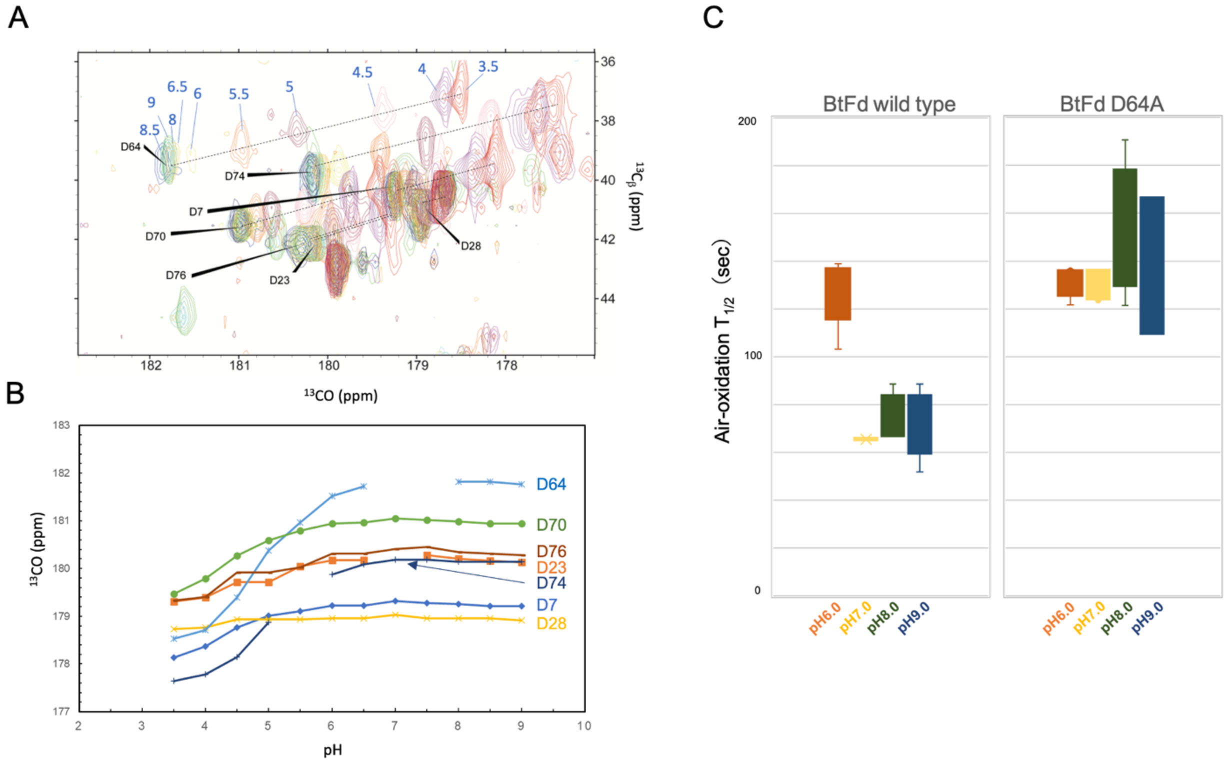


**Fig. S7. The p*K*a estimation of the Asp64 in *Bt*Fd** (A) A close-up view of an overlay of the of the CACO experiments (Cβ offset) of NMR. The assignments were indicated. For Asp64 signals, corresponding pH values are shown in blue as a representative of titrations. Chemical shift changes are indicated by black dotted line for each residue. Detail information were described in the supplemental method and text. (B)Titration curves obtained from the CACO experiments (Cβ offset). The points correspond to pH 7.0 and 7.5 are missing in a series of Asp64, and the point corresponds to pH 6.5 is missing in a series of Asp74. (C) The oxidation rates of *Bt*Fd wild-type and D64A under the different pH conditions.

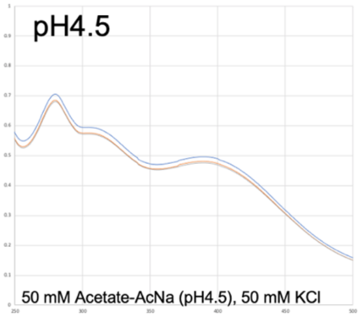


| | 0h | 15h | 25h |
|---|---|---|---|
| 304nm/280nm | 0.842 | 0.841 | 0.840 |
| 388nm/280nm | 0.702 | 0.702 | 0.699 |
| 304nm/388nm | 1.199 | 1.198 | 1.202 |

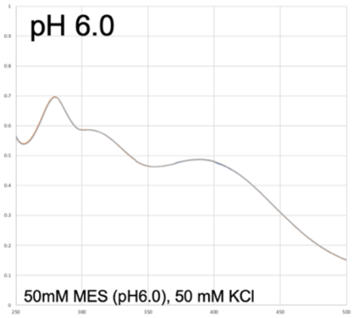


| | 0h | 15h | 25h |
|---|---|---|---|
| 304nm/280nm | 0.842 | 0.842 | 0.842 |
| 388nm/280nm | 0.700 | 0.700 | 0.701 |
| 304nm/388nm | 1.204 | 1.203 | 1.201 |

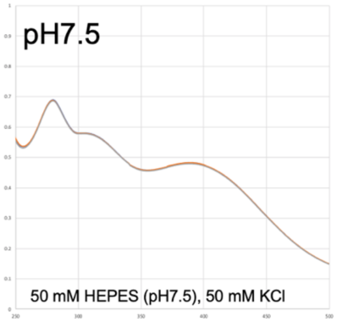


| | 0h | 15h | 25h |
|---|---|---|---|
| 304nm/280nm | 0.842 | 0.842 | 0.842 |
| 388nm/280nm | 0.700 | 0.702 | 0.698 |
| 304nm/388nm | 1.201 | 1.200 | 1.206 |

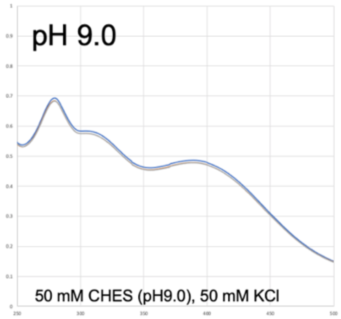


| | 0h | 15h | 25h |
|---|---|---|---|
| 304nm/280nm | 0.842 | 0.842 | 0.842 |
| 388nm/280nm | 0.702 | 0.700 | 0.700 |
| 304nm/388nm | 1.200 | 1.202 | 1.203 |

**Fig. S8. Stability of [4Fe-4S] cluster in *Bt*Fd under the different pH conditions.** The UV-Vis absorbance ratios of 304 nm/280 nm, 388 nm/280 nm and 304 nm/388 nm are shown in the table below each spectral panel.

**Table S1.** Cartesian coordinate of $CM_H$ used for the DFT calculations

| atom | *x* / Å | *y* / Å | *z* / Å | *x* / Å | *y* / Å | *z* / Å | *x* / Å |
|---|---|---|---|---|---|---|---|
| Fe | 51.156 | 26.907 | -3.179 | C | 54.743 | 28.460 | -1.565 |
| Fe | 48.980 | 28.449 | -3.585 | H | 56.554 | 27.390 | 0.793 |
| Fe | 50.443 | 27.558 | -5.708 | H | 55.043 | 29.582 | 1.547 |
| Fe | 51.472 | 29.467 | -3.999 | H | 56.424 | 29.637 | 0.416 |
| S | 49.681 | 29.696 | -5.349 | H | 53.459 | 29.555 | -0.186 |
| S | 52.523 | 27.621 | -4.827 | H | 54.736 | 30.548 | -0.942 |
| S | 50.558 | 28.775 | -2.049 | H | 55.562 | 28.730 | -2.232 |
| S | 49.293 | 26.313 | -4.232 | N | 53.408 | 26.740 | 1.466 |
| N | 48.058 | 30.492 | -0.277 | C | 52.479 | 26.302 | 2.502 |
| C | 48.219 | 31.919 | -0.009 | C | 52.033 | 24.858 | 2.334 |
| C | 48.373 | 32.739 | -1.294 | O | 51.037 | 24.440 | 2.919 |
| O | 48.790 | 33.898 | -1.240 | C | 51.274 | 27.224 | 2.544 |
| C | 49.362 | 32.179 | 0.994 | O | 50.717 | 27.319 | 1.226 |
| C | 50.663 | 31.594 | 0.458 | C | 51.683 | 28.622 | 3.012 |
| C | 49.044 | 31.580 | 2.341 | H | 52.994 | 26.365 | 3.461 |
| C | 51.905 | 32.039 | 1.231 | H | 50.532 | 26.832 | 3.239 |
| H | 47.297 | 32.268 | 0.456 | H | 50.801 | 29.255 | 3.109 |
| H | 49.470 | 33.257 | 1.113 | H | 52.182 | 28.559 | 3.980 |
| H | 50.608 | 30.506 | 0.515 | H | 52.365 | 29.072 | 2.291 |
| H | 50.788 | 31.907 | -0.579 | H | 52.996 | 26.764 | 0.533 |
| H | 49.851 | 31.818 | 3.035 | H | 49.739 | 27.288 | 1.273 |
| H | 48.106 | 32.000 | 2.704 | N | 52.739 | 24.097 | 1.524 |
| H | 48.952 | 30.499 | 2.237 | C | 52.447 | 22.665 | 1.388 |
| H | 51.935 | 31.514 | 2.186 | C | 50.999 | 22.421 | 0.957 |
| H | 52.792 | 31.797 | 0.647 | O | 50.377 | 21.434 | 1.348 |
| H | 51.851 | 33.114 | 1.399 | C | 52.807 | 21.865 | 2.656 |
| H | 48.919 | 30.035 | -0.577 | C | 54.316 | 21.837 | 2.927 |
| N | 48.006 | 32.177 | -2.459 | O | 55.107 | 21.756 | 1.965 |
| C | 47.922 | 32.940 | -3.709 | H | 53.083 | 22.274 | 0.594 |
| C | 49.249 | 33.579 | -4.101 | H | 52.318 | 22.322 | 3.516 |
| O | 49.289 | 34.735 | -4.534 | H | 52.466 | 20.836 | 2.537 |
| C | 46.811 | 33.991 | -3.658 | H | 53.515 | 24.428 | 0.950 |
| H | 47.015 | 34.684 | -2.842 | N | 50.463 | 23.310 | 0.125 |
| H | 46.787 | 34.530 | -4.605 | C | 48.604 | 24.612 | -0.759 |
| H | 45.856 | 33.492 | -3.491 | O | 48.641 | 25.501 | 0.378 |
| H | 47.761 | 31.192 | -2.562 | H | 49.257 | 24.999 | -1.542 |
| N | 50.339 | 32.821 | -3.970 | H | 47.581 | 24.541 | -1.129 |
| C | 51.682 | 33.345 | -4.188 | H | 50.971 | 24.109 | -0.253 |
| C | 52.133 | 33.338 | -5.642 | H | 49.569 | 25.652 | 0.655 |
| O | 53.113 | 34.018 | -5.958 | H | 53.842 | 28.258 | -2.144 |
| C | 52.691 | 32.574 | -3.340 | C | 54.867 | 25.648 | -2.573 |

| | | | | | | | |
|---|---|---|---|---|---|---|---|
| S | 53.161 | 30.980 | -3.986 | H | 55.881 | 25.386 | -2.790 |
| H | 51.680 | 34.387 | -3.867 | H | 54.491 | 26.286 | -3.345 |
| H | 53.599 | 33.172 | -3.255 | C | 49.084 | 23.239 | -0.339 |
| H | 52.258 | 32.412 | -2.353 | H | 48.350 | 22.946 | 0.382 |
| H | 50.323 | 31.834 | -3.713 | H | 49.166 | 22.483 | -1.092 |
| N | 51.471 | 32.597 | -6.531 | O | 54.717 | 21.901 | 4.118 |
| C | 51.828 | 32.597 | -7.941 | H | 55.079 | 21.052 | 4.381 |
| C | 52.971 | 31.678 | -8.370 | C | 44.136 | 29.698 | -6.554 |
| O | 53.291 | 31.646 | -9.569 | O | 43.373 | 29.594 | -7.527 |
| H | 50.948 | 32.303 | -8.513 | C | 42.720 | 30.361 | -4.590 |
| H | 52.110 | 33.612 | -8.220 | C | 42.430 | 31.428 | -3.521 |
| H | 50.685 | 31.990 | -6.300 | C | 42.981 | 29.008 | -3.942 |
| N | 53.583 | 30.912 | -7.458 | C | 43.569 | 31.718 | -2.601 |
| C | 55.472 | 29.647 | -6.552 | H | 41.838 | 30.289 | -5.226 |
| H | 55.963 | 30.501 | -6.085 | H | 42.169 | 32.359 | -4.023 |
| H | 54.745 | 29.214 | -5.865 | H | 41.594 | 31.087 | -2.910 |
| H | 56.214 | 28.897 | -6.829 | H | 43.199 | 28.278 | -4.721 |
| H | 53.297 | 30.825 | -6.483 | H | 43.831 | 29.097 | -3.266 |
| C | 46.889 | 29.856 | -0.139 | H | 42.094 | 28.705 | -3.386 |
| H | 47.678 | 32.225 | -4.495 | H | 43.174 | 31.997 | -1.624 |
| S | 46.811 | 29.076 | -3.190 | H | 44.189 | 30.826 | -2.510 |
| C | 46.231 | 28.098 | -1.820 | H | 44.157 | 32.539 | -3.012 |
| H | 45.445 | 28.621 | -1.316 | N | 45.177 | 28.897 | -6.365 |
| H | 45.862 | 27.161 | -2.182 | C | 46.957 | 27.364 | -7.072 |
| O | 45.859 | 30.407 | 0.236 | H | 47.143 | 26.500 | -7.710 |
| C | 46.900 | 28.370 | -0.456 | H | 47.689 | 28.145 | -7.280 |
| H | 46.342 | 27.619 | 0.062 | H | 47.015 | 27.070 | -6.024 |
| H | 47.940 | 28.260 | -0.231 | H | 45.775 | 28.909 | -5.539 |
| C | 54.749 | 30.117 | -7.808 | C | 45.558 | 27.903 | -7.361 |
| H | 55.572 | 30.565 | -8.324 | H | 45.577 | 28.326 | -8.344 |
| H | 54.248 | 29.433 | -8.461 | H | 44.800 | 27.150 | -7.311 |
| C | 55.189 | 26.023 | -1.124 | C | 43.907 | 30.756 | -5.480 |
| O | 55.475 | 25.135 | -0.325 | H | 44.808 | 30.834 | -4.908 |
| C | 53.647 | 24.744 | -2.630 | H | 43.682 | 31.704 | -5.922 |
| S | 52.213 | 25.438 | -1.764 | C | 51.979 | 26.168 | -8.183 |
| H | 53.369 | 24.590 | -3.673 | S | 50.437 | 27.038 | -7.918 |
| H | 53.892 | 23.789 | -2.165 | H | 51.743 | 25.253 | -8.726 |
| N | 55.102 | 27.304 | -0.742 | H | 52.385 | 25.925 | -7.201 |
| C | 55.518 | 27.685 | 0.626 | C | 53.058 | 26.931 | -8.974 |
| C | 54.676 | 27.075 | 1.726 | H | 52.720 | 27.928 | -9.167 |
| O | 55.181 | 26.899 | 2.841 | H | 53.962 | 26.965 | -8.403 |
| C | 55.433 | 29.219 | 0.596 | H | 53.241 | 26.430 | -9.902 |
| C | 54.492 | 29.567 | -0.536 | | | | |

**Table S2.** Examined charge/spin states with $CM_{NA}$ model for (A) oxidized and (B) reduced states. Definition of Fe1–Fe4 is illustrated using CM below. o12 and r5 (indicated by yellow highlighted) were most stable states for *Ox* and *Red* states, respectively.

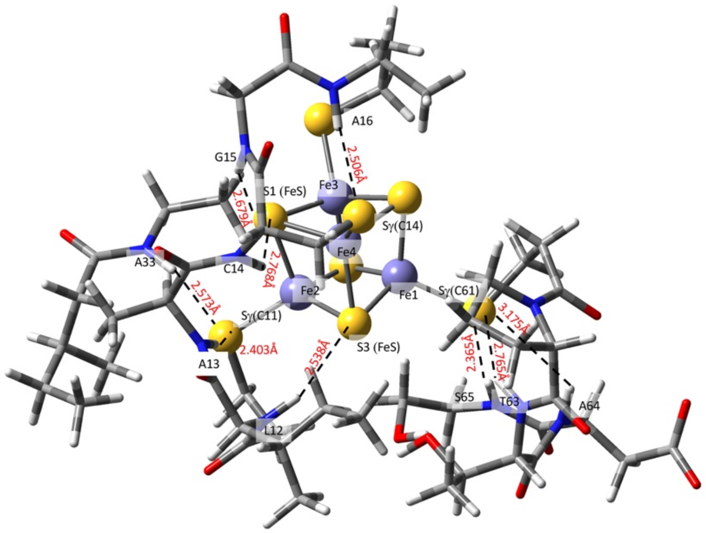


**(A) Oxidized (*Ox*) state (u and d represent up and down spins)**

| Examined states | Fe1 | Fe2 | Fe3 | Fe4 | Energy / Hartree |
|---|---|---|---|---|---|
| charge | **2** | **2** | **3** | **3** | |
| o1 | u | d | u | d | -8400.022791 |
| o2 | u | d | d | u | -8400.025020 |
| charge | **2** | **3** | **2** | **3** | |
| o3 | u | u | d | d | -8399.983854 |
| o4 | u | d | d | u | -8400.025019 |
| charge | **2** | **3** | **3** | **2** | |
| o5 | u | u | d | d | -8400.022702 |
| o6 | u | d | u | d | -8400.022789 |
| charge | **3** | **2** | **3** | **2** | |
| o7 | u | u | d | d | -8400.022689 |
| o8 | d | u | u | d | -8400.025019 |
| charge | **3** | **2** | **2** | **3** | |
| o9 | u | u | d | d | -8400.022693 |
| o10 | d | u | d | u | -8400.022801 |
| charge | **3** | **3** | **2** | **2** | |
| o11 | u | d | u | d | -8400.022802 |
| o12 | d | u | u | d | -8400.025022 |

**(B) Reduced (*Red*) state (u and d represent up and down spins)**

| Examined states | Fe1 | Fe2 | Fe3 | Fe4 | Energy / Hartree |
|---|---|---|---|---|---|
| charge | 3 | 2 | 2 | 2 | |
| r1 | u | u | d | d | -8399.871923 |
| r2 | u | d | u | d | -8399.870871 |
| r3 | u | d | d | u | -8399.871694 |
| charge | 2 | 3 | 2 | 2 | |
| r4 | u | u | d | d | -8399.871876 |
| r5 | d | u | u | d | -8399.872594 |
| r6 | d | u | d | u | -8399.869132 |
| charge | 2 | 2 | 3 | 2 | |
| r7 | u | d | u | d | -8399.870871 |
| r8 | d | u | u | d | -8399.872589 |
| r9 | d | d | u | u | -8399.867054 |
| charge | 2 | 2 | 2 | 3 | |
| r10 | u | d | d | u | -8399.871693 |
| r11 | d | u | d | u | -8399.869131 |
| r12 | d | d | u | u | -8399.656590 |

**Table S3.** Cartesian coordinate (in Å) of before and after the geometry optimization. (The Cartesian coordinate before the geometry optimization is the neutron structure)

Part 1. **CM** model

| Before optimization | | | | After optimization | | | |
|---|---|---|---|---|---|---|---|
| Fe | 51.1560 | 26.9070 | -3.1790 | Fe | 51.0899 | 26.8241 | -3.0748 |
| Fe | 48.9800 | 28.4490 | -3.5850 | Fe | 48.9077 | 28.4118 | -3.6890 |
| Fe | 50.4430 | 27.5580 | -5.7080 | Fe | 50.4026 | 27.5837 | -5.8198 |
| Fe | 51.4720 | 29.4670 | -3.9990 | Fe | 51.5173 | 29.5281 | -4.0769 |
| S | 49.6810 | 29.6960 | -5.3490 | S | 49.6463 | 29.7914 | -5.5251 |
| S | 52.5230 | 27.6210 | -4.8270 | S | 52.5245 | 27.5200 | -4.9853 |
| S | 50.5580 | 28.7750 | -2.0490 | S | 50.4549 | 28.9082 | -2.0530 |
| S | 49.2930 | 26.3130 | -4.2320 | S | 49.2273 | 26.1675 | -4.3358 |
| N | 48.0580 | 30.4920 | -0.2770 | N | 48.0580 | 30.4920 | -0.2770 |
| C | 48.2190 | 31.9190 | -0.0090 | C | 48.2190 | 31.9190 | -0.0090 |
| C | 48.3730 | 32.7390 | -1.2940 | C | 48.3730 | 32.7390 | -1.2940 |
| O | 48.7900 | 33.8980 | -1.2400 | O | 48.7900 | 33.8980 | -1.2400 |
| C | 49.3620 | 32.1790 | 0.9940 | C | 49.3620 | 32.1790 | 0.9940 |
| C | 50.6630 | 31.5940 | 0.4580 | C | 50.6630 | 31.5940 | 0.4580 |
| C | 49.0440 | 31.5800 | 2.3410 | C | 49.0440 | 31.5800 | 2.3410 |
| C | 51.9050 | 32.0390 | 1.2310 | C | 51.9050 | 32.0390 | 1.2310 |
| H | 47.2970 | 32.2680 | 0.4560 | H | 47.2970 | 32.2680 | 0.4560 |
| H | 49.4700 | 33.2570 | 1.1130 | H | 49.4700 | 33.2570 | 1.1130 |
| H | 50.6080 | 30.5060 | 0.5150 | H | 50.6080 | 30.5060 | 0.5150 |
| H | 50.7880 | 31.9070 | -0.5790 | H | 50.7880 | 31.9070 | -0.5790 |
| H | 49.8510 | 31.8180 | 3.0350 | H | 49.8510 | 31.8180 | 3.0350 |
| H | 48.1060 | 32.0000 | 2.7040 | H | 48.1060 | 32.0000 | 2.7040 |
| H | 48.9520 | 30.4990 | 2.2370 | H | 48.9520 | 30.4990 | 2.2370 |
| H | 51.9350 | 31.5140 | 2.1860 | H | 51.9350 | 31.5140 | 2.1860 |
| H | 52.7920 | 31.7970 | 0.6470 | H | 52.7920 | 31.7970 | 0.6470 |
| H | 51.8510 | 33.1140 | 1.3990 | H | 51.8510 | 33.1140 | 1.3990 |
| H | 48.9190 | 30.0350 | -0.5770 | H | 48.9190 | 30.0350 | -0.5770 |
| N | 48.0060 | 32.1770 | -2.4590 | N | 48.0060 | 32.1770 | -2.4590 |
| C | 47.9220 | 32.9400 | -3.7090 | C | 47.9220 | 32.9400 | -3.7090 |

| | | | | | | | |
|---|---|---|---|---|---|---|---|
| C | 49.2490 | 33.5790 | -4.1010 | C | 49.2490 | 33.5790 | -4.1010 |
| O | 49.2890 | 34.7350 | -4.5340 | O | 49.2890 | 34.7350 | -4.5340 |
| C | 46.8110 | 33.9910 | -3.6580 | C | 46.8110 | 33.9910 | -3.6580 |
| H | 47.0150 | 34.6840 | -2.8420 | H | 47.0150 | 34.6840 | -2.8420 |
| H | 46.7870 | 34.5300 | -4.6050 | H | 46.7870 | 34.5300 | -4.6050 |
| H | 45.8560 | 33.4920 | -3.4910 | H | 45.8560 | 33.4920 | -3.4910 |
| H | 47.7610 | 31.1920 | -2.5620 | H | 47.7610 | 31.1920 | -2.5620 |
| N | 50.3390 | 32.8210 | -3.9700 | N | 50.3390 | 32.8210 | -3.9700 |
| C | 51.6820 | 33.3450 | -4.1880 | C | 51.6821 | 33.3451 | -4.1880 |
| C | 52.1330 | 33.3380 | -5.6420 | C | 52.1330 | 33.3380 | -5.6420 |
| O | 53.1130 | 34.0180 | -5.9580 | O | 53.1130 | 34.0180 | -5.9580 |
| C | 52.6910 | 32.5740 | -3.3400 | C | 52.6908 | 32.5738 | -3.3400 |
| S | 53.1610 | 30.9800 | -3.9860 | S | 53.2923 | 30.9518 | -3.9777 |
| H | 51.6800 | 34.3870 | -3.8670 | H | 51.6800 | 34.3870 | -3.8670 |
| H | 53.5990 | 33.1720 | -3.2550 | H | 53.5990 | 33.1720 | -3.2550 |
| H | 52.2580 | 32.4120 | -2.3530 | H | 52.2581 | 32.4121 | -2.3530 |
| H | 50.3230 | 31.8340 | -3.7130 | H | 50.3230 | 31.8340 | -3.7130 |
| N | 51.4710 | 32.5970 | -6.5310 | N | 51.4710 | 32.5970 | -6.5310 |
| C | 51.8280 | 32.5970 | -7.9410 | C | 51.8280 | 32.5970 | -7.9410 |
| C | 52.9710 | 31.6780 | -8.3700 | C | 52.9710 | 31.6780 | -8.3700 |
| O | 53.2910 | 31.6460 | -9.5690 | O | 53.2910 | 31.6460 | -9.5690 |
| H | 50.9480 | 32.3030 | -8.5130 | H | 50.9480 | 32.3030 | -8.5130 |
| H | 52.1100 | 33.6120 | -8.2200 | H | 52.1100 | 33.6120 | -8.2200 |
| H | 50.6850 | 31.9900 | -6.3000 | H | 50.6850 | 31.9900 | -6.3000 |
| N | 53.5830 | 30.9120 | -7.4580 | N | 53.5830 | 30.9120 | -7.4580 |
| C | 55.4720 | 29.6470 | -6.5520 | C | 55.4720 | 29.6470 | -6.5520 |
| H | 55.9630 | 30.5010 | -6.0850 | H | 55.9630 | 30.5010 | -6.0850 |
| H | 54.7450 | 29.2140 | -5.8650 | H | 54.7450 | 29.2140 | -5.8650 |
| H | 56.2140 | 28.8970 | -6.8290 | H | 56.2140 | 28.8970 | -6.8290 |
| H | 53.2970 | 30.8250 | -6.4830 | H | 53.2970 | 30.8250 | -6.4830 |
| C | 46.8895 | 29.8555 | -0.1395 | C | 46.8895 | 29.8555 | -0.1395 |
| H | 47.6780 | 32.2250 | -4.4950 | H | 47.6780 | 32.2250 | -4.4950 |

| | | | | | | | |
|---|---|---|---|---|---|---|---|
| S | 46.8110 | 29.0760 | -3.1900 | S | 46.7360 | 29.1131 | -3.2770 |
| C | 46.2315 | 28.0983 | -1.8201 | C | 46.2318 | 28.0982 | -1.8202 |
| H | 45.4454 | 28.6209 | -1.3162 | H | 45.4454 | 28.6209 | -1.3162 |
| H | 45.8624 | 27.1613 | -2.1815 | H | 45.8623 | 27.1613 | -2.1815 |
| O | 45.8590 | 30.4070 | 0.2360 | O | 45.8590 | 30.4070 | 0.2360 |
| C | 46.9000 | 28.3700 | -0.4560 | C | 46.8999 | 28.3701 | -0.4560 |
| H | 46.3416 | 27.6187 | 0.0623 | H | 46.3415 | 27.6187 | 0.0623 |
| H | 47.9403 | 28.2600 | -0.2310 | H | 47.9403 | 28.2599 | -0.2310 |
| C | 54.7490 | 30.1170 | -7.8080 | C | 54.7490 | 30.1170 | -7.8080 |
| H | 55.5724 | 30.5645 | -8.3243 | H | 55.5724 | 30.5645 | -8.3243 |
| H | 54.2480 | 29.4329 | -8.4607 | H | 54.2480 | 29.4329 | -8.4607 |
| C | 55.1890 | 26.0230 | -1.1240 | C | 55.1890 | 26.0230 | -1.1240 |
| O | 55.4750 | 25.1350 | -0.3250 | O | 55.4750 | 25.1350 | -0.3250 |
| C | 53.6470 | 24.7440 | -2.6300 | C | 53.6470 | 24.7441 | -2.6301 |
| S | 52.2130 | 25.4380 | -1.7640 | S | 52.2470 | 25.4672 | -1.6975 |
| H | 53.3690 | 24.5900 | -3.6730 | H | 53.3690 | 24.5900 | -3.6730 |
| H | 53.8920 | 23.7890 | -2.1650 | H | 53.8920 | 23.7890 | -2.1650 |
| N | 55.1020 | 27.3040 | -0.7420 | N | 55.1020 | 27.3040 | -0.7420 |
| C | 55.5180 | 27.6850 | 0.6260 | C | 55.5180 | 27.6850 | 0.6260 |
| C | 54.6760 | 27.0750 | 1.7260 | C | 54.6760 | 27.0750 | 1.7260 |
| O | 55.1810 | 26.8990 | 2.8410 | O | 55.1810 | 26.8990 | 2.8410 |
| C | 55.4330 | 29.2190 | 0.5960 | C | 55.4330 | 29.2190 | 0.5960 |
| C | 54.4920 | 29.5670 | -0.5360 | C | 54.4920 | 29.5670 | -0.5360 |
| C | 54.7430 | 28.4600 | -1.5650 | C | 54.7430 | 28.4600 | -1.5650 |
| H | 56.5540 | 27.3900 | 0.7930 | H | 56.5540 | 27.3900 | 0.7930 |
| H | 55.0430 | 29.5820 | 1.5470 | H | 55.0430 | 29.5820 | 1.5470 |
| H | 56.4240 | 29.6370 | 0.4160 | H | 56.4240 | 29.6370 | 0.4160 |
| H | 53.4590 | 29.5550 | -0.1860 | H | 53.4590 | 29.5550 | -0.1860 |
| H | 54.7360 | 30.5480 | -0.9420 | H | 54.7360 | 30.5480 | -0.9420 |
| H | 55.5620 | 28.7300 | -2.2320 | H | 55.5620 | 28.7300 | -2.2320 |
| N | 53.4080 | 26.7400 | 1.4660 | N | 53.4080 | 26.7400 | 1.4660 |
| C | 52.4790 | 26.3020 | 2.5020 | C | 52.4790 | 26.3020 | 2.5020 |

| | | | | | | | |
|---|---|---|---|---|---|---|---|
| C | 52.0330 | 24.8580 | 2.3340 | C | 52.0330 | 24.8580 | 2.3340 |
| O | 51.0370 | 24.4400 | 2.9190 | O | 51.0370 | 24.4400 | 2.9190 |
| C | 51.2740 | 27.2240 | 2.5440 | C | 51.2740 | 27.2240 | 2.5440 |
| O | 50.7170 | 27.3190 | 1.2260 | O | 50.7170 | 27.3190 | 1.2260 |
| C | 51.6830 | 28.6220 | 3.0120 | C | 51.6830 | 28.6220 | 3.0120 |
| H | 52.9940 | 26.3650 | 3.4610 | H | 52.9940 | 26.3650 | 3.4610 |
| H | 50.5320 | 26.8320 | 3.2390 | H | 50.5320 | 26.8320 | 3.2390 |
| H | 50.8010 | 29.2550 | 3.1090 | H | 50.8010 | 29.2550 | 3.1090 |
| H | 52.1820 | 28.5590 | 3.9800 | H | 52.1820 | 28.5590 | 3.9800 |
| H | 52.3650 | 29.0720 | 2.2910 | H | 52.3650 | 29.0720 | 2.2910 |
| H | 52.9960 | 26.7640 | 0.5330 | H | 52.9960 | 26.7640 | 0.5330 |
| H | 49.7390 | 27.2880 | 1.2730 | H | 49.7390 | 27.2880 | 1.2730 |
| N | 52.7390 | 24.0970 | 1.5240 | N | 52.7390 | 24.0970 | 1.5240 |
| C | 52.4470 | 22.6650 | 1.3880 | C | 52.4470 | 22.6650 | 1.3880 |
| C | 50.9990 | 22.4210 | 0.9570 | C | 50.9990 | 22.4210 | 0.9570 |
| O | 50.3770 | 21.4340 | 1.3480 | O | 50.3770 | 21.4340 | 1.3480 |
| C | 52.8070 | 21.8650 | 2.6560 | C | 52.8070 | 21.8650 | 2.6560 |
| C | 54.3160 | 21.8370 | 2.9270 | C | 54.3160 | 21.8370 | 2.9270 |
| O | 55.1070 | 21.7560 | 1.9650 | O | 55.1070 | 21.7560 | 1.9650 |
| O | 54.7150 | 21.9010 | 4.1110 | O | 54.7150 | 21.9010 | 4.1110 |
| H | 53.0830 | 22.2740 | 0.5940 | H | 53.0830 | 22.2740 | 0.5940 |
| H | 52.3180 | 22.3220 | 3.5160 | H | 52.3180 | 22.3220 | 3.5160 |
| H | 52.4660 | 20.8360 | 2.5370 | H | 52.4660 | 20.8360 | 2.5370 |
| H | 53.5150 | 24.4280 | 0.9500 | H | 53.5150 | 24.4280 | 0.9500 |
| N | 50.4630 | 23.3100 | 0.1250 | N | 50.4630 | 23.3100 | 0.1250 |
| C | 48.6040 | 24.6120 | -0.7590 | C | 48.6040 | 24.6120 | -0.7590 |
| O | 48.6410 | 25.5010 | 0.3780 | O | 48.6410 | 25.5010 | 0.3780 |
| H | 49.2570 | 24.9990 | -1.5420 | H | 49.2570 | 24.9990 | -1.5420 |
| H | 47.5810 | 24.5410 | -1.1290 | H | 47.5810 | 24.5410 | -1.1290 |
| H | 50.9710 | 24.1090 | -0.2530 | H | 50.9710 | 24.1090 | -0.2530 |
| H | 49.5690 | 25.6520 | 0.6550 | H | 49.5690 | 25.6520 | 0.6550 |
| H | 53.8420 | 28.2580 | -2.1440 | H | 53.8420 | 28.2580 | -2.1440 |

| | | | | | | | |
|---|---|---|---|---|---|---|---|
| C | 54.8670 | 25.6480 | -2.5730 | C | 54.8670 | 25.6480 | -2.5730 |
| H | 55.8815 | 25.3859 | -2.7899 | H | 55.8815 | 25.3859 | -2.7899 |
| H | 54.4906 | 26.2860 | -3.3451 | H | 54.4906 | 26.2860 | -3.3451 |
| C | 49.0840 | 23.2390 | -0.3390 | C | 49.0840 | 23.2390 | -0.3390 |
| H | 48.3495 | 22.9463 | 0.3820 | H | 48.3495 | 22.9463 | 0.3820 |
| H | 49.1657 | 22.4831 | -1.0919 | H | 49.1657 | 22.4831 | -1.0919 |
| C | 44.1360 | 29.6980 | -6.5540 | C | 44.1360 | 29.6980 | -6.5540 |
| O | 43.3730 | 29.5940 | -7.5270 | O | 43.3730 | 29.5940 | -7.5270 |
| C | 42.7200 | 30.3610 | -4.5900 | C | 42.7200 | 30.3610 | -4.5900 |
| C | 42.4300 | 31.4280 | -3.5210 | C | 42.4300 | 31.4280 | -3.5210 |
| C | 42.9810 | 29.0080 | -3.9420 | C | 42.9810 | 29.0080 | -3.9420 |
| C | 43.5690 | 31.7180 | -2.6010 | C | 43.5690 | 31.7180 | -2.6010 |
| H | 41.8380 | 30.2890 | -5.2260 | H | 41.8380 | 30.2890 | -5.2260 |
| H | 42.1690 | 32.3590 | -4.0230 | H | 42.1690 | 32.3590 | -4.0230 |
| H | 41.5940 | 31.0870 | -2.9100 | H | 41.5940 | 31.0870 | -2.9100 |
| H | 43.1990 | 28.2780 | -4.7210 | H | 43.1990 | 28.2780 | -4.7210 |
| H | 43.8310 | 29.0970 | -3.2660 | H | 43.8310 | 29.0970 | -3.2660 |
| H | 42.0940 | 28.7050 | -3.3860 | H | 42.0940 | 28.7050 | -3.3860 |
| H | 43.1740 | 31.9970 | -1.6240 | H | 43.1740 | 31.9970 | -1.6240 |
| H | 44.1890 | 30.8260 | -2.5100 | H | 44.1890 | 30.8260 | -2.5100 |
| H | 44.1570 | 32.5390 | -3.0120 | H | 44.1570 | 32.5390 | -3.0120 |
| N | 45.1770 | 28.8970 | -6.3650 | N | 45.1770 | 28.8970 | -6.3650 |
| C | 46.9570 | 27.3640 | -7.0720 | C | 46.9570 | 27.3640 | -7.0720 |
| H | 47.1430 | 26.5000 | -7.7100 | H | 47.1430 | 26.5000 | -7.7100 |
| H | 47.6890 | 28.1450 | -7.2800 | H | 47.6890 | 28.1450 | -7.2800 |
| H | 47.0150 | 27.0700 | -6.0240 | H | 47.0150 | 27.0700 | -6.0240 |
| H | 45.7750 | 28.9090 | -5.5390 | H | 45.7750 | 28.9090 | -5.5390 |
| C | 45.5580 | 27.9030 | -7.3610 | C | 45.5580 | 27.9030 | -7.3610 |
| H | 45.5770 | 28.3258 | -8.3437 | H | 45.5770 | 28.3258 | -8.3437 |
| H | 44.7997 | 27.1498 | -7.3114 | H | 44.7997 | 27.1498 | -7.3114 |
| C | 43.9070 | 30.7560 | -5.4800 | C | 43.9070 | 30.7560 | -5.4800 |
| H | 44.8080 | 30.8336 | -4.9080 | H | 44.8080 | 30.8336 | -4.9080 |

| | | | | | | | |
|---|---|---|---|---|---|---|---|
| H | 43.6824 | 31.7042 | -5.9221 | H | 43.6824 | 31.7042 | -5.9221 |
| C | 51.9790 | 26.1680 | -8.1830 | C | 51.9790 | 26.1682 | -8.1826 |
| S | 50.4370 | 27.0380 | -7.9180 | S | 50.3589 | 27.0411 | -8.0554 |
| H | 51.7430 | 25.2530 | -8.7260 | H | 51.7430 | 25.2530 | -8.7261 |
| H | 52.3850 | 25.9250 | -7.2010 | H | 52.3850 | 25.9249 | -7.2010 |
| C | 53.0578 | 26.9308 | -8.9741 | C | 53.0578 | 26.9307 | -8.9743 |
| H | 52.7204 | 27.9276 | -9.1674 | H | 52.7204 | 27.9276 | -9.1673 |
| H | 53.9619 | 26.9647 | -8.4028 | H | 53.9619 | 26.9647 | -8.4029 |
| H | 53.2407 | 26.4300 | -9.9018 | H | 53.2407 | 26.4300 | -9.9019 |

Part 2. **$CM_H$** model

| Before optimization | | | | After optimization | | | |
|---|---|---|---|---|---|---|---|
| Fe | 51.1560 | 26.9070 | -3.1790 | Fe | 51.0943 | 26.8246 | -3.0613 |
| Fe | 48.9800 | 28.4489 | -3.5851 | Fe | 48.9111 | 28.3918 | -3.6611 |
| Fe | 50.4430 | 27.5580 | -5.7080 | Fe | 50.4148 | 27.5566 | -5.7999 |
| Fe | 51.4720 | 29.4670 | -3.9990 | Fe | 51.5223 | 29.5247 | -4.0565 |
| S | 49.6810 | 29.6960 | -5.3490 | S | 49.6625 | 29.7604 | -5.4967 |
| S | 52.5230 | 27.6210 | -4.8270 | S | 52.5387 | 27.4961 | -4.9493 |
| S | 50.5580 | 28.7750 | -2.0490 | S | 50.4679 | 28.8805 | -2.0194 |
| S | 49.2930 | 26.3130 | -4.2320 | S | 49.2421 | 26.1403 | -4.3041 |
| N | 48.0580 | 30.4920 | -0.2770 | N | 48.0580 | 30.4920 | -0.2770 |
| C | 48.2190 | 31.9190 | -0.0090 | C | 48.2190 | 31.9190 | -0.0090 |
| C | 48.3730 | 32.7390 | -1.2940 | C | 48.3730 | 32.7390 | -1.2940 |
| O | 48.7900 | 33.8980 | -1.2400 | O | 48.7900 | 33.8980 | -1.2400 |
| C | 49.3620 | 32.1790 | 0.9940 | C | 49.3620 | 32.1790 | 0.9940 |
| C | 50.6630 | 31.5940 | 0.4580 | C | 50.6630 | 31.5940 | 0.4580 |
| C | 49.0440 | 31.5800 | 2.3410 | C | 49.0440 | 31.5800 | 2.3410 |
| C | 51.9050 | 32.0390 | 1.2310 | C | 51.9050 | 32.0390 | 1.2310 |
| H | 47.2970 | 32.2680 | 0.4560 | H | 47.2970 | 32.2680 | 0.4560 |
| H | 49.4700 | 33.2570 | 1.1130 | H | 49.4700 | 33.2570 | 1.1130 |

| | | | | | | | |
|---|---|---|---|---|---|---|---|
| H | 50.6080 | 30.5060 | 0.5150 | H | 50.6080 | 30.5060 | 0.5150 |
| H | 50.7880 | 31.9070 | -0.5790 | H | 50.7880 | 31.9070 | -0.5790 |
| H | 49.8510 | 31.8180 | 3.0350 | H | 49.8510 | 31.8180 | 3.0350 |
| H | 48.1060 | 32.0000 | 2.7040 | H | 48.1060 | 32.0000 | 2.7040 |
| H | 48.9520 | 30.4990 | 2.2370 | H | 48.9520 | 30.4990 | 2.2370 |
| H | 51.9350 | 31.5140 | 2.1860 | H | 51.9350 | 31.5140 | 2.1860 |
| H | 52.7920 | 31.7970 | 0.6470 | H | 52.7920 | 31.7970 | 0.6470 |
| H | 51.8510 | 33.1140 | 1.3990 | H | 51.8510 | 33.1140 | 1.3990 |
| H | 48.9190 | 30.0350 | -0.5770 | H | 48.9190 | 30.0350 | -0.5770 |
| N | 48.0060 | 32.1770 | -2.4590 | N | 48.0060 | 32.1770 | -2.4590 |
| C | 47.9220 | 32.9400 | -3.7090 | C | 47.9220 | 32.9400 | -3.7090 |
| C | 49.2490 | 33.5790 | -4.1010 | C | 49.2490 | 33.5790 | -4.1010 |
| O | 49.2890 | 34.7350 | -4.5340 | O | 49.2890 | 34.7350 | -4.5340 |
| C | 46.8110 | 33.9910 | -3.6580 | C | 46.8110 | 33.9910 | -3.6580 |
| H | 47.0150 | 34.6840 | -2.8420 | H | 47.0150 | 34.6840 | -2.8420 |
| H | 46.7870 | 34.5300 | -4.6050 | H | 46.7870 | 34.5300 | -4.6050 |
| H | 45.8560 | 33.4920 | -3.4910 | H | 45.8560 | 33.4920 | -3.4910 |
| H | 47.7610 | 31.1920 | -2.5620 | H | 47.7610 | 31.1920 | -2.5620 |
| N | 50.3390 | 32.8210 | -3.9700 | N | 50.3390 | 32.8210 | -3.9700 |
| C | 51.6820 | 33.3450 | -4.1880 | C | 51.6820 | 33.3451 | -4.1880 |
| C | 52.1330 | 33.3380 | -5.6420 | C | 52.1330 | 33.3380 | -5.6420 |
| O | 53.1130 | 34.0180 | -5.9580 | O | 53.1130 | 34.0180 | -5.9580 |
| C | 52.6910 | 32.5740 | -3.3400 | C | 52.6908 | 32.5738 | -3.3400 |
| S | 53.1610 | 30.9800 | -3.9860 | S | 53.2990 | 30.9481 | -3.9670 |
| H | 51.6800 | 34.3870 | -3.8670 | H | 51.6800 | 34.3870 | -3.8670 |
| H | 53.5990 | 33.1720 | -3.2550 | H | 53.5990 | 33.1720 | -3.2550 |
| H | 52.2580 | 32.4120 | -2.3530 | H | 52.2581 | 32.4121 | -2.3530 |
| H | 50.3230 | 31.8340 | -3.7130 | H | 50.3230 | 31.8340 | -3.7130 |
| N | 51.4710 | 32.5970 | -6.5310 | N | 51.4710 | 32.5970 | -6.5310 |
| C | 51.8280 | 32.5970 | -7.9410 | C | 51.8280 | 32.5970 | -7.9410 |
| C | 52.9710 | 31.6780 | -8.3700 | C | 52.9710 | 31.6780 | -8.3700 |
| O | 53.2910 | 31.6460 | -9.5690 | O | 53.2910 | 31.6460 | -9.5690 |

| | | | | | | | |
|---|---|---|---|---|---|---|---|
| H | 50.9480 | 32.3030 | -8.5130 | H | 50.9480 | 32.3030 | -8.5130 |
| H | 52.1100 | 33.6120 | -8.2200 | H | 52.1100 | 33.6120 | -8.2200 |
| H | 50.6850 | 31.9900 | -6.3000 | H | 50.6850 | 31.9900 | -6.3000 |
| N | 53.5830 | 30.9120 | -7.4580 | N | 53.5830 | 30.9120 | -7.4580 |
| C | 55.4720 | 29.6470 | -6.5520 | C | 55.4720 | 29.6470 | -6.5520 |
| H | 55.9630 | 30.5010 | -6.0850 | H | 55.9630 | 30.5010 | -6.0850 |
| H | 54.7450 | 29.2140 | -5.8650 | H | 54.7450 | 29.2140 | -5.8650 |
| H | 56.2140 | 28.8970 | -6.8290 | H | 56.2140 | 28.8970 | -6.8290 |
| H | 53.2970 | 30.8250 | -6.4830 | H | 53.2970 | 30.8250 | -6.4830 |
| C | 46.8895 | 29.8555 | -0.1395 | C | 46.8895 | 29.8555 | -0.1395 |
| H | 47.6780 | 32.2250 | -4.4950 | H | 47.6780 | 32.2250 | -4.4950 |
| S | 46.8110 | 29.0760 | -3.1900 | S | 46.7475 | 29.1063 | -3.2748 |
| C | 46.2315 | 28.0983 | -1.8201 | C | 46.2318 | 28.0982 | -1.8202 |
| H | 45.4454 | 28.6209 | -1.3162 | H | 45.4454 | 28.6209 | -1.3163 |
| H | 45.8624 | 27.1613 | -2.1815 | H | 45.8623 | 27.1613 | -2.1815 |
| O | 45.8590 | 30.4070 | 0.2360 | O | 45.8590 | 30.4070 | 0.2360 |
| C | 46.9000 | 28.3700 | -0.4560 | C | 46.8999 | 28.3701 | -0.4560 |
| H | 46.3416 | 27.6187 | 0.0623 | H | 46.3415 | 27.6187 | 0.0623 |
| H | 47.9403 | 28.2600 | -0.2310 | H | 47.9403 | 28.2599 | -0.2310 |
| C | 54.7490 | 30.1170 | -7.8080 | C | 54.7490 | 30.1170 | -7.8080 |
| H | 55.5724 | 30.5645 | -8.3243 | H | 55.5724 | 30.5645 | -8.3244 |
| H | 54.2481 | 29.4329 | -8.4607 | H | 54.2481 | 29.4329 | -8.4607 |
| C | 55.1890 | 26.0230 | -1.1240 | C | 55.1890 | 26.0230 | -1.1240 |
| O | 55.4750 | 25.1350 | -0.3250 | O | 55.4750 | 25.1350 | -0.3250 |
| C | 53.6470 | 24.7440 | -2.6300 | C | 53.6469 | 24.7441 | -2.6301 |
| S | 52.2130 | 25.4380 | -1.7640 | S | 52.2521 | 25.4563 | -1.6763 |
| H | 53.3690 | 24.5900 | -3.6730 | H | 53.3690 | 24.5900 | -3.6730 |
| H | 53.8920 | 23.7890 | -2.1650 | H | 53.8920 | 23.7890 | -2.1650 |
| N | 55.1020 | 27.3040 | -0.7420 | N | 55.1020 | 27.3040 | -0.7420 |
| C | 55.5180 | 27.6850 | 0.6260 | C | 55.5180 | 27.6850 | 0.6260 |
| C | 54.6760 | 27.0750 | 1.7260 | C | 54.6760 | 27.0750 | 1.7260 |
| O | 55.1810 | 26.8990 | 2.8410 | O | 55.1810 | 26.8990 | 2.8410 |

| | | | | | | | |
|---|---|---|---|---|---|---|---|
| C | 55.4330 | 29.2190 | 0.5960 | C | 55.4330 | 29.2190 | 0.5960 |
| C | 54.4920 | 29.5670 | -0.5360 | C | 54.4920 | 29.5670 | -0.5360 |
| C | 54.7430 | 28.4600 | -1.5650 | C | 54.7430 | 28.4600 | -1.5650 |
| H | 56.5540 | 27.3900 | 0.7930 | H | 56.5540 | 27.3900 | 0.7930 |
| H | 55.0430 | 29.5820 | 1.5470 | H | 55.0430 | 29.5820 | 1.5470 |
| H | 56.4240 | 29.6370 | 0.4160 | H | 56.4240 | 29.6370 | 0.4160 |
| H | 53.4590 | 29.5550 | -0.1860 | H | 53.4590 | 29.5550 | -0.1860 |
| H | 54.7360 | 30.5480 | -0.9420 | H | 54.7360 | 30.5480 | -0.9420 |
| H | 55.5620 | 28.7300 | -2.2320 | H | 55.5620 | 28.7300 | -2.2320 |
| N | 53.4080 | 26.7400 | 1.4660 | N | 53.4080 | 26.7400 | 1.4660 |
| C | 52.4790 | 26.3020 | 2.5020 | C | 52.4790 | 26.3020 | 2.5020 |
| C | 52.0330 | 24.8580 | 2.3340 | C | 52.0330 | 24.8580 | 2.3340 |
| O | 51.0370 | 24.4400 | 2.9190 | O | 51.0370 | 24.4400 | 2.9190 |
| C | 51.2740 | 27.2240 | 2.5440 | C | 51.2740 | 27.2240 | 2.5440 |
| O | 50.7170 | 27.3190 | 1.2260 | O | 50.7170 | 27.3190 | 1.2260 |
| C | 51.6830 | 28.6220 | 3.0120 | C | 51.6830 | 28.6220 | 3.0120 |
| H | 52.9940 | 26.3650 | 3.4610 | H | 52.9940 | 26.3650 | 3.4610 |
| H | 50.5320 | 26.8320 | 3.2390 | H | 50.5320 | 26.8320 | 3.2390 |
| H | 50.8010 | 29.2550 | 3.1090 | H | 50.8010 | 29.2550 | 3.1090 |
| H | 52.1820 | 28.5590 | 3.9800 | H | 52.1820 | 28.5590 | 3.9800 |
| H | 52.3650 | 29.0720 | 2.2910 | H | 52.3650 | 29.0720 | 2.2910 |
| H | 52.9960 | 26.7640 | 0.5330 | H | 52.9960 | 26.7640 | 0.5330 |
| H | 49.7390 | 27.2880 | 1.2730 | H | 49.7390 | 27.2880 | 1.2730 |
| N | 52.7390 | 24.0970 | 1.5240 | N | 52.7390 | 24.0970 | 1.5240 |
| C | 52.4470 | 22.6650 | 1.3880 | C | 52.4470 | 22.6650 | 1.3880 |
| C | 50.9990 | 22.4210 | 0.9570 | C | 50.9990 | 22.4210 | 0.9570 |
| O | 50.3770 | 21.4340 | 1.3480 | O | 50.3770 | 21.4340 | 1.3480 |
| C | 52.8070 | 21.8650 | 2.6560 | C | 52.8070 | 21.8650 | 2.6560 |
| C | 54.3160 | 21.8370 | 2.9270 | C | 54.3160 | 21.8370 | 2.9270 |
| O | 55.1070 | 21.7560 | 1.9650 | O | 55.1070 | 21.7560 | 1.9650 |
| H | 53.0830 | 22.2740 | 0.5940 | H | 53.0830 | 22.2740 | 0.5940 |
| H | 52.3180 | 22.3220 | 3.5160 | H | 52.3180 | 22.3220 | 3.5160 |

| | | | | | | | |
|---|---|---|---|---|---|---|---|
| H | 52.4660 | 20.8360 | 2.5370 | H | 52.4660 | 20.8360 | 2.5370 |
| H | 53.5150 | 24.4280 | 0.9500 | H | 53.5150 | 24.4280 | 0.9500 |
| N | 50.4630 | 23.3100 | 0.1250 | N | 50.4630 | 23.3100 | 0.1250 |
| C | 48.6040 | 24.6120 | -0.7590 | C | 48.6040 | 24.6120 | -0.7590 |
| O | 48.6410 | 25.5010 | 0.3780 | O | 48.6410 | 25.5010 | 0.3780 |
| H | 49.2570 | 24.9990 | -1.5420 | H | 49.2570 | 24.9990 | -1.5420 |
| H | 47.5810 | 24.5410 | -1.1290 | H | 47.5810 | 24.5410 | -1.1290 |
| H | 50.9710 | 24.1090 | -0.2530 | H | 50.9710 | 24.1090 | -0.2530 |
| H | 49.5690 | 25.6520 | 0.6550 | H | 49.5690 | 25.6520 | 0.6550 |
| H | 53.8420 | 28.2580 | -2.1440 | H | 53.8420 | 28.2580 | -2.1440 |
| C | 54.8670 | 25.6480 | -2.5730 | C | 54.8670 | 25.6480 | -2.5730 |
| H | 55.8815 | 25.3859 | -2.7899 | H | 55.8815 | 25.3859 | -2.7899 |
| H | 54.4906 | 26.2860 | -3.3451 | H | 54.4906 | 26.2860 | -3.3451 |
| C | 49.0840 | 23.2390 | -0.3390 | C | 49.0840 | 23.2390 | -0.3390 |
| H | 48.3495 | 22.9463 | 0.3820 | H | 48.3495 | 22.9463 | 0.3820 |
| H | 49.1657 | 22.4831 | -1.0919 | H | 49.1657 | 22.4831 | -1.0919 |
| O | 54.7173 | 21.9014 | 4.1179 | O | 54.7173 | 21.9014 | 4.1179 |
| H | 55.6962 | 21.9043 | 4.0821 | H | 55.6962 | 21.9043 | 4.0821 |
| C | 44.1360 | 29.6980 | -6.5540 | C | 44.1360 | 29.6980 | -6.5540 |
| O | 43.3730 | 29.5940 | -7.5270 | O | 43.3730 | 29.5940 | -7.5270 |
| C | 42.7200 | 30.3610 | -4.5900 | C | 42.7200 | 30.3610 | -4.5900 |
| C | 42.4300 | 31.4280 | -3.5210 | C | 42.4300 | 31.4280 | -3.5210 |
| C | 42.9810 | 29.0080 | -3.9420 | C | 42.9810 | 29.0080 | -3.9420 |
| C | 43.5690 | 31.7180 | -2.6010 | C | 43.5690 | 31.7180 | -2.6010 |
| H | 41.8380 | 30.2890 | -5.2260 | H | 41.8380 | 30.2890 | -5.2260 |
| H | 42.1690 | 32.3590 | -4.0230 | H | 42.1690 | 32.3590 | -4.0230 |
| H | 41.5940 | 31.0870 | -2.9100 | H | 41.5940 | 31.0870 | -2.9100 |
| H | 43.1990 | 28.2780 | -4.7210 | H | 43.1990 | 28.2780 | -4.7210 |
| H | 43.8310 | 29.0970 | -3.2660 | H | 43.8310 | 29.0970 | -3.2660 |
| H | 42.0940 | 28.7050 | -3.3860 | H | 42.0940 | 28.7050 | -3.3860 |
| H | 43.1740 | 31.9970 | -1.6240 | H | 43.1740 | 31.9970 | -1.6240 |
| H | 44.1890 | 30.8260 | -2.5100 | H | 44.1890 | 30.8260 | -2.5100 |

| | | | | | | | |
|---|---|---|---|---|---|---|---|
| H | 44.1570 | 32.5390 | -3.0120 | H | 44.1570 | 32.5390 | -3.0120 |
| N | 45.1770 | 28.8970 | -6.3650 | N | 45.1770 | 28.8970 | -6.3650 |
| C | 46.9570 | 27.3640 | -7.0720 | C | 46.9570 | 27.3640 | -7.0720 |
| H | 47.1430 | 26.5000 | -7.7100 | H | 47.1430 | 26.5000 | -7.7100 |
| H | 47.6890 | 28.1450 | -7.2800 | H | 47.6890 | 28.1450 | -7.2800 |
| H | 47.0150 | 27.0700 | -6.0240 | H | 47.0150 | 27.0700 | -6.0240 |
| H | 45.7750 | 28.9090 | -5.5390 | H | 45.7750 | 28.9090 | -5.5390 |
| C | 45.5580 | 27.9030 | -7.3610 | C | 45.5580 | 27.9030 | -7.3610 |
| H | 45.5771 | 28.3258 | -8.3437 | H | 45.5771 | 28.3258 | -8.3437 |
| H | 44.7997 | 27.1498 | -7.3114 | H | 44.7997 | 27.1498 | -7.3114 |
| C | 43.9070 | 30.7560 | -5.4800 | C | 43.9070 | 30.7560 | -5.4800 |
| H | 44.8080 | 30.8336 | -4.9080 | H | 44.8080 | 30.8336 | -4.9080 |
| H | 43.6824 | 31.7042 | -5.9221 | H | 43.6824 | 31.7042 | -5.9221 |
| C | 51.9790 | 26.1680 | -8.1830 | C | 51.9790 | 26.1682 | -8.1827 |
| S | 50.4370 | 27.0380 | -7.9180 | S | 50.3626 | 27.0425 | -8.0337 |
| H | 51.7430 | 25.2530 | -8.7260 | H | 51.7430 | 25.2530 | -8.7260 |
| H | 52.3850 | 25.9250 | -7.2010 | H | 52.3850 | 25.9249 | -7.2010 |
| C | 53.0578 | 26.9308 | -8.9741 | C | 53.0578 | 26.9307 | -8.9743 |
| H | 52.7204 | 27.9276 | -9.1674 | H | 52.7204 | 27.9276 | -9.1674 |
| H | 53.9619 | 26.9647 | -8.4028 | H | 53.9619 | 26.9647 | -8.4029 |
| H | 53.2408 | 26.4300 | -9.9018 | H | 53.2407 | 26.4300 | -9.9019 |

**Table S4.** Calculated IP values of the $CM_H$ and CM models with and without the environment effect.

| | IP(red) values | |
|---|---|---|
| | Gas phase | IEFPCM (Eps=4) (*7*) |
| **$CM_H$** | –1.06 | 2.59 |
| **CM** | –3.27 | 1.62 |
| ΔIP | 2.21 | 0.97 |

**Table S5. The distance between the acidic residues and the [4Fe-4S] cluster.***

| | | Distance between the Cγ of Asp and the [4Fe-4S] cluster, (Å) | Remarks |
|---|---|---|---|
| D7 | | 10.5 | |
| D23 | | 16.6 | |
| D26 | | 12.0 | |
| D28 | | 11.3 | |
| D30 | | 13.7 | |
| D38 | | 16.2 | |
| D39 | | 16.9 | |
| D48 | | 24.0 | |
| D52 | | 19.2 | |
| D53 | | 15.1 | |
| D56 | | 13.7 | |
| D64 | | 10.1 | |
| D70 | | 23.0 | |
| D74 | | 22.4 | |
| D76 | | 18.2 | |

| | | Distance(s) between the Cδ of Glu and the [4Fe-4S] cluster, (Å) | Remarks |
|---|---|---|---|
| E9 | | 13.0 | |
| E45 | | 22.4<br>21.7 | Alternative conformations |
| E59 | | 12.8 | |
| E71 | | 23.4<br>22.9<br>22.2 | Alternative conformations |
| E81 | | 16.4 | |

* Distances are shown by measuring the center-of-gravity coordinates of the [4Fe-4S] cluster and the sidechain terminal carbon of acidic residues.
